# MacroH2A impedes metastatic growth by enforcing a discrete dormancy program in disseminated cancer cells

**DOI:** 10.1101/2021.12.07.471619

**Authors:** Dan Sun, Dan Filipescu, Dan Hasson, Deepak K. Singh, Saul Carcamo, Bassem Khalil, Brett A. Miles, William Westra, Karl Christoph Sproll, Emily Bernstein, Julio A. Aguirre-Ghiso

**Author notes:** Department of Cell Biology, Cancer Dormancy and Tumor Microenvironment Institute, Gruss-Lipper Biophotonics Center, Albert Einstein Cancer Center, Albert Einstein College of Medicine, 1300 Morris Park Avenue, Bronx, NY. USA. Western Atlantic University School of Medicine, Plantation, FL.

## Abstract

MacroH2A variants have been associated with tumor suppression through inhibition of proliferation and metastasis, as well as their role in cellular senescence. However, their role in regulating the dormant state of disseminated cancer cells (DCCs) remains unclear. Here we reveal that solitary dormant DCCs display increased levels of macroH2A variants in head and neck squamous cell carcinoma PDX models and patient samples compared to proliferating primary or metastatic lesions. We further demonstrate that microenvironmental and stress adaptive signals such as TGFβ2 and p38α/β, which induce DCC dormancy, upregulate macroH2A expression. Functionally, we find that overexpression of macroH2A variants is sufficient to induce tumor cells into dormancy and notably, inducible expression of the macroH2A2 variant suppresses the growth of DCCs into overt metastasis. However, this dormant state does not require well-characterized dormancy factors such as DEC2 and NR2F1, suggesting alternate pathways. Our transcriptomic analyses reveal that macroH2A2 overexpression inhibits E2F, RAS and MYC signaling programs, while upregulating inflammatory cytokines commonly secreted by senescent cells. Taken together, our results demonstrate that macroH2A2 enforces a stable dormant phenotype in DCCs by activating a select subset of dormancy and senescence genes that limit metastasis initiation.

## Introduction

Microenvironmental and cellular intrinsic mechanisms control the quiescence of disseminated cancer cells (DCCs) during dormancy, a stage of cancer cell quiescence^1-3^ that can precede lethal metastasis of solid cancers by many years^4^. However, less is known about the epigenetic mechanisms controlling dormancy in cancer cells with genetically altered genomes^5^. In cancer models and human specimens, specific microenvironmental cues such as TGFβ2^6^, retinoic acid^7^, BMP7^8^, periostin^9^, LIF^10^ and GAS6^11^ can activate transcriptional programs of quiescence, pluripotency, and survival in dormant tumor cells. We initially reported that in HNSCC and breast cancer models, a low ratio of the extracellular signal-regulated kinase (ERK)/p38α/β kinase activities can predict a state of cancer cell dormancy *in vivo* ^12, 13^ and regulates a specific dormancy gene expression signature^6, 14^. One of the major transcription factors (TFs) of the dormancy program is NR2F1, a nuclear orphan receptor that contributes to a repressive chromatin state in dormant cancer cells characterized by high levels of H3K27me3 and H3K9me3^7, 15^. Combined treatment of 5-azacytidine and retinoid acid can induce dormancy in malignant cells and is also associated with the induction of a global chromatin repressive state^7^. Similarly, a deep growth arrest that is activated in senescent cells is associated with mechanisms of chromatin repression and induction of senescence associated heterochromatin foci (SAHF)^16-18^. Thus, persistent growth arrest states such as dormancy and senescence exhibit transcriptional and chromatin alterations.

While epigenetic mechanisms control tumor onset, progression and relapse^19^ and altered chromatin states can modulate the cell cycle and cellular responses to the microenvironment^17, 20^, our knowledge of the epigenetic mechanisms guided by histone variants in the regulation of DCC fate remains limited. The H2A family of histones contains multiple variants, of which macroH2A is unique in having a large and evolutionarily conserved macro domain at the C-terminus. In mammals, two different genes, *MACROH2A1* and *MACROH2A2*, encode macroH2A1 and macroH2A2, respectively, and macroH2A1.1 and macroH2A1.2 are alternatively spliced variants encoded by *MACROH2A1*. MacroH2A variants are generally considered transcriptionally repressive histones that co-localize genome-wide with histone modifications such as H3K27me3 and H3K9me3. This includes the inactive X chromosome in mammalian female cells^21^ and broad repressive regions across autosomal chromatin^21^, as well as SAHF^16^. In somatic cells, these histone variants function as barriers in the reprogramming of differentiated cells towards pluripotency^22-24^. Therefore, macroH2A variants may limit cellular plasticity in differentiated cells.

In keeping with this concept, macroH2A expression is lost or attenuated at different stages of cancer progression across multiple tumor types^19, 25-27^, and is generally considered tumor suppressive. For example, the abundance of macroH2A correlates inversely with proliferation in a lung cancer recurrence study^27^. Moreover, the transcriptional loss of macroH2A correlates with melanoma metastasis in patients, while its overexpression significantly suppresses lung metastasis in mice^28^. However, the mechanisms governed by macroH2A in inhibiting metastasis initiation remain unclear. Here we show that the histone variant macroH2A plays a significant role in the induction of a unique program of DCC dormancy that suppresses metastasis. Using *in vitro* and *in vivo* models, as well as validation in human patient HNSCC DCCs, we show that macroH2A variants, and particularly macroH2A2, potently suppress metastasis. This occurs via a transcriptional program that involves inhibition of RAS and MYC signaling, and induction of cytokines that are produced by senescent cells. Thus, macroH2A variants suppress metastasis via a dormancy program that contains components of the senescence program. This could be potentially exploited to force DCCs into a deep growth arrest and prevent metastasis initiation.

## Results

### MacroH2A1 expression is enriched in dormant vs. proliferative HNSCC cells and DCCs

Taking advantage of HNSCC PDX models T-HEp3 (proliferative) and D-HEp3 (dormant) cells, which display these respective phenotypes when inoculated *in vivo* in the chick embryo chorioallantoic membrane (CAM) or nude mice (**Supplementary Fig.1A**)^6, 12, 29^, we examined expression of macroH2A1 and macroH2A2. By performing combined RNA-seq and epigenomic analyses, we found that macroH2A1 levels are higher in D-HEp3 compared to T-HEp3 cells *in vitro* (**Supplementary Fig.1B**), and the macroH2A1 promoter is marked by H3K27ac. In contrast, macroH2A2 is not expressed in either proliferative or dormant HEp3 variants at the mRNA or protein levels, and its locus is coated by the repressive histone modification H3K27me3 (**Supplementary Fig.1B-C**). As reported^30^, D-HEp3 (but not T-HEp3) cells inoculated on CAMs synchronously enter a sustained G0/G1 arrest forming small fully quiescent tumor micro-masses. Analysis of macroH2A1 protein levels via immunofluorescence (IF) directly from the CAM revealed that it was significantly upregulated in D-HEp3 vs. T-HEp3 cells (**Fig.1A-B**). The mRNA level of macroH2A1 is also enriched in D-HEp3 over T-HEp3 cells in CAMs, along with dormancy-associated transcription factors (TFs), NR2F1 and DEC2 (**Fig.1C**). Given that increased expression of NR2F1 and DEC2 in D-HEp3 cells is dependent on p38 α/β signaling^6, 7^, we treated D-HEp3 cells with a p38 α/β inhibitor (SB203580) and observed a ∼50% reduction in macroH2A1 mRNA levels (**Fig.1D, Supplementary Fig.1D**). We have also shown that DEC2 and p38α/β activation in D-HEp3 cells is dependent on TGFβ2^6, 31^. Upon daily treatments of TGFβ2 for 6 days, the growth of T-HEp3 cells *in vivo* was efficiently inhibited (**Supplementary Fig.1E**) as reported^6, 31^, which significantly increased macroH2A1 mRNA levels (**Fig.1E**). MacroH2A2 expression remained undetectable even after TGFβ2 treatment (data not shown). Therefore, macroH2A1 is positively regulated by the TGFβ2 and p38α/β pathways in HEp3 cells and is associated with a dormancy program.

**Figure 1.**
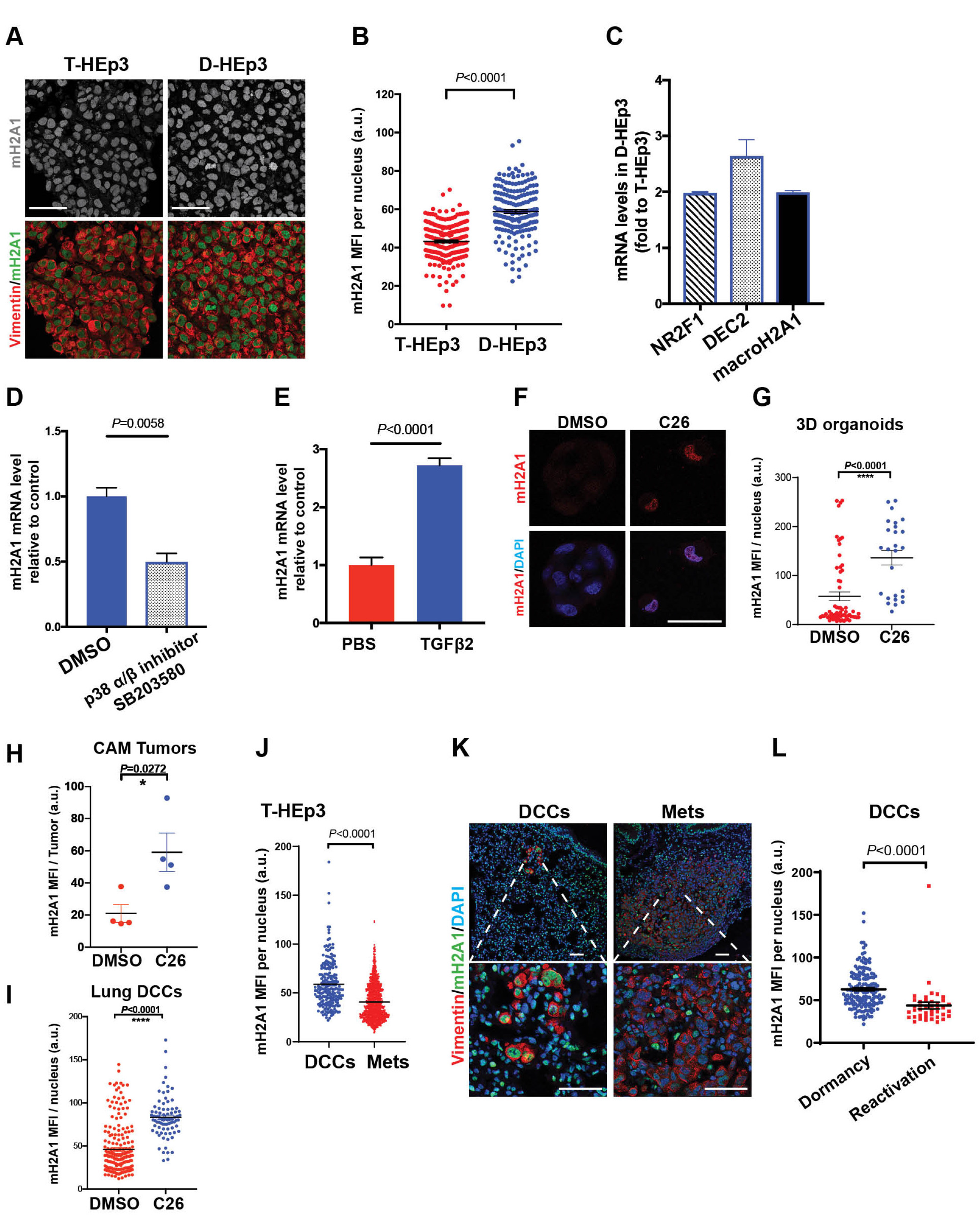
MacroH2A1 expression is enriched in dormant vs. proliferative HNSCC cells and DCCs. (A-B) Images and quantification of T-HEp3 or D-HEp3 CAM tumors stained for macroH2A1 (green). Tumor sections were also stained with Vimentin (red) and DAPI (blue). Representative images were shown. Cells (n>100) were assessed in each group, Mann-Whitney test. Scale bars, 50um. (C) Relative mRNA expression levels in D-HEp3 cells, compared to that in T-HEp3 cells. (D) D-HEp3 cells were treated with DMSO or SB 203580 (5uM, 48hrs) in serum free medium *in vitro*. See Supplementary Fig.1D for the drug efficacy. Q-PCR measured the relative mRNA transcript level of macroH2A1 to control/ DMSO. PCR in triplicate, mean + s.e.m., two-tailed Student’s t-test. (E) qPCR-measured macroH2A1 mRNA levels in T-HEp3 CAM tumors treated with or without 10ng/mL TGFβ2 for 6 days. T-HEp3 cells were pre-treated with 10ng/mL TGFβ2 for 24hr in serum-free medium *in vitro* before CAM inoculation. n=4 tumors per group, PCR in triplicate, mean + s.e.m., two-tailed Student’s t-test. (F-G) Images and quantification of T-HEp3 cells grown in 3D organoids and cells were treated with 0.2uM C26 (NR2F1 agonist) or DMSO as indicated for 4 days *in vitro*. Cells were stained for macroH2A1 (green) and DAPI (blue). Scale bars, 25um. (H-I) Bar graphs show the quantification of macroH2A1 MFI in T-HEp3 cells grown in CAM model (H) or DCCs found in mouse lungs (I). mean + s.e.m., two-tailed Student’s t-test. (J-L) Experimental metastasis assay. T-HEp3 cells were tail vein injected in nude mice. One or three weeks later, the lungs were retrieved and processed for FFPE sections. See Supplementary Fig.1F for the assay scheme. n = 10 animals were assessed. (J) Quantification of macroH2A1 mean fluorescent intensity (MFI) per nucleus in either DCCs or metastatic lesions. Singles cells and small clusters (<20 cells) are all included as DCC events. Mann-Whitney test. (K) IF representative images show the stain for macroH2A1(green), human vimentin (red) and DAPI (blue). Scale bars, 50um. (L) Quantification of macroH2A1 MFI in DCC events found in 1 week after tail-vein injection (Dormancy) or 3 weeks after tail-vein injection (Reactivation). n=165 cells pooled from 5 animals at Dormancy phase, n=41 cells pooled from 5 animals at Reactivation phase. Unpaired Student’s t-test.

We also explored whether NR2F1, which can be induced by morphogens like *all-trans* retinoic acid (atRA) and BMP7 to induce dormancy, could upregulate macroH2A1 expression. To this end we made use of a recently identified NR2F1 agonist (C26)^32^, which induces a dormancy-like program and suppresses metastasis in the HEp3 HNSCC model^32^. T-HEp3 cells were treated with C26 in 3D organoid cultures for four days, which caused a robust upregulation of macroH2A1 protein levels (**Fig.1F-G**). By treating T-HEp3 cells *in vivo* in the CAM, we also found that growth suppression induced by C26 was associated with a strong induction of macroH2A1 in these dormant-like lesions (**Fig.1H**). Finally, because C26 prevents solitary DCCs from forming proliferative metastasis, we tested whether this dormant solitary DCC state was associated with macroH2A induction. In accordance, C26-mediated suppression of metastasis in lungs^32^ was associated with an upregulation of macroH2A1 in DCCs (**Fig.1I**), which also upregulate NR2F1 and p27, while downregulating H3S10 phosphorylation levels^32^. We conclude that NR2F1 activity may directly or indirectly regulate macroH2A1.

We next tested whether macroH2A expression is different between metastatic solitary DCCs vs. proliferative lesions in secondary organs, as solitary DCCs can be found in a non-proliferative dormant state^15, 32-35^. T-HEp3 cells, which can efficiently colonize distal organs in both spontaneous and experimental metastasis models^6, 36^ were tail vein injected in nude mice. DCCs that arrive in the lung in either of these assays can persist in a dormant state for at least six weeks, while a fraction reactivate and resume proliferation as early as two weeks after arrival to the lung^16, 29, 37, 38^. We harvested lungs at one or three weeks after i.v. injection to detect dormant DCCs at high frequency at week 1 (dormancy) and reactivating DCCs at 3 weeks (reactivation) (**Supplementary Fig.1F**). To locate human solitary DCCs, DCC clusters and metastasis in murine lungs, we used an antibody against the intermediate filament vimentin, which preferentially binds to the human over mouse antigen^6, 7, 36^ and co-stained for macroH2A1. We considered single cells, doublets, and clusters of less than 20-cells (previously found to be devoid of proliferation markers^6, 7, 36^) as dormant DCC events; events with >20 cells were categorized as proliferative metastasis, as they commonly express proliferation markers^6, 7, 36^. When comparing by cluster size, we found that macroH2A1 was significantly higher in solitary and small DCC cluster populations compared to metastases (**Fig.1J-K**). When focusing only on solitary and small DCC cluster events (<20 cell/cluster) during dormancy (week 1) and reactivation phases (week 3), we found that the macroH2A1 signal was higher in DCCs in dormancy (**Fig.1L**). This data suggests a potential role for macroH2A in enforcing a dormant state of DCCs. Collectively, the above experiments support that signaling and transcriptional mechanisms that control cancer cell dormancy downstream of microenvironmental cues can promote macroH2A expression.

### MacroH2A variants are enriched in solitary DCCs vs. primary tumor and metastatic lesions in HNSCC patients

We next tested whether the association of macroH2A1 and macroH2A2 expression in DCCs held true in HNSCC patients by quantifying macroH2A variant expression in FFPE samples from poorly differentiated tumors. Examination of seven pairs of primary tumors (MS-PT) and matching lymph node metastases (MS-LN Met) *via* IF revealed that both macroH2A1 and macroH2A2 signals decrease in LN metastases vs. primary tumors (**Fig. 2A-B, Supplementary Fig.2A-D**). While the T-HEp3 PDX model and SQ20B cells do not express macroH2A2, patient samples, albeit with great heterogeneity, and the FaDu cell line do express macroH2A2. Thus, macroH2A2 silencing seems to occur heterogeneously. In addition, we observed heterogeneity in macroH2A expression patterns in both MS-PT and MS-LN Met samples across patients. Even within one patient, macroH2A expression is heterogenous in different regions of primary tumors and lymph node metastasis, suggesting dynamic regulation of its expression. Nevertheless, careful single cell quantification revealed that the proportion of macroH2A1^High^ cells was significantly reduced in 57% (4/7) of patient lymph node metastases compared to primary lesions (**Supplementary Fig.2A**). Strikingly, the percentage of macroH2A2^High^ cells significantly decreased in 86% (6/7) of patient LN metastases compared to primary lesions (**Supplementary Fig.2B**).

**Figure 2.**
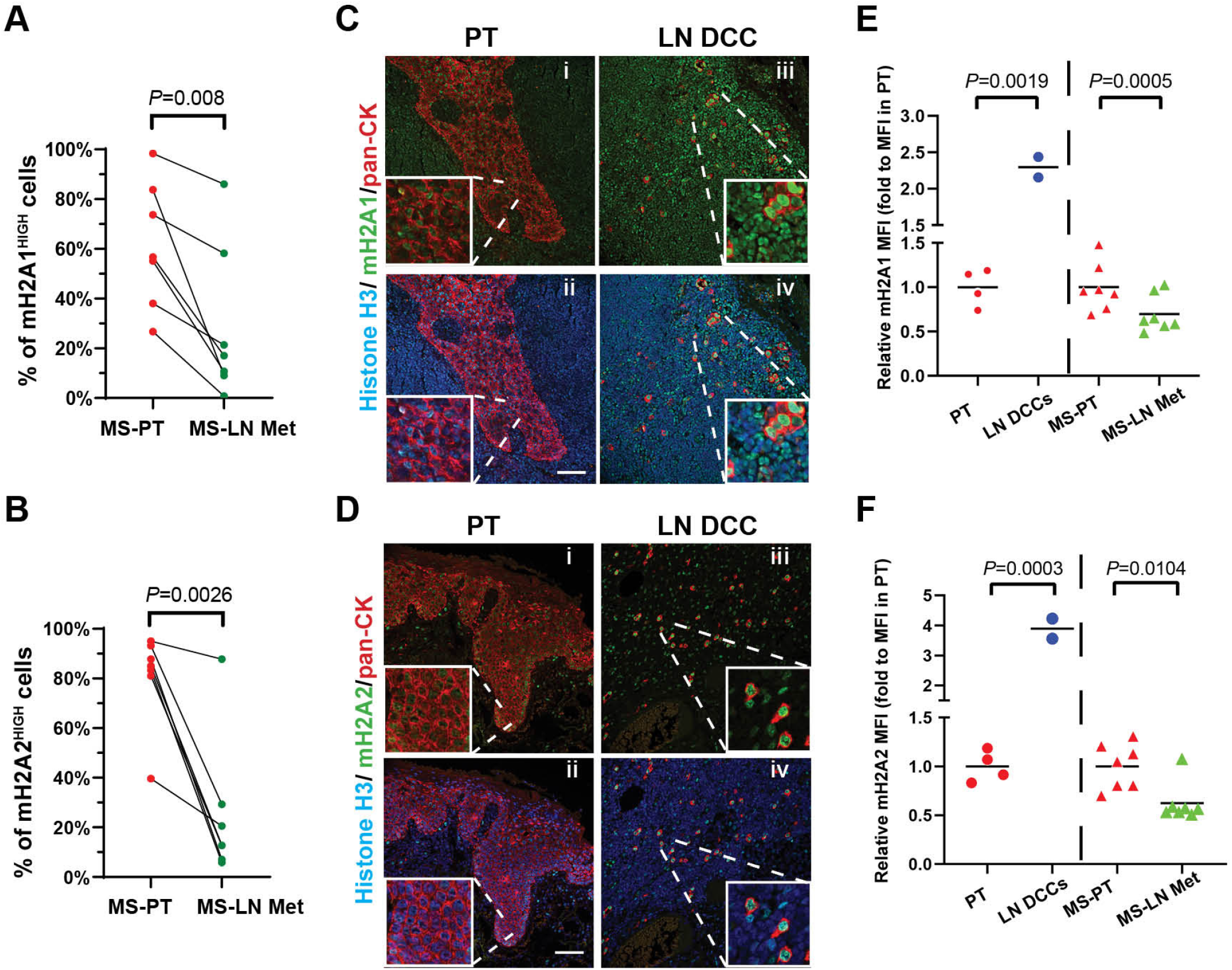
MacroH2A variants are enriched in solitary DCCs vs. primary tumor and metastatic lesions in HNSCC patients. (A-B) Quantification of the percentage of macroH2A1 or macroH2A2 high cells in either primary tumors or lymph node metastases. Each paired primary tumor (MS-PT) tissue section (red) and Lymph Node Metastatic lesion (MS-LN Met, green) were from the same patient. See Supplemental Fig.2 for the detail quantification method and representative images. n=7 patients; each data point represents the mean percentage of macroH2A ^HIGH^ cells pooled from 5∼6 fields of view. Paired Student’s t-test. (C-D) Representative images of IF staining in primary tumor tissues or clinically negative LN tissues. macroH2A1(C) or macroH2A2(D) in green, pan-cytokeratin in red and HistoneH3 in blue. Scale bar, 75um. (E-F) Measurement of relative macroH2A1 or macroH2A2 intensity (normalized to Histone H3 intensity) in HNSCC primary tumors (red dots) or DCCs (blue dots) found in clinically negative patient LN tissues after the primary tumor removal, as well as paired PT section (red triangles) and Lymph Node Metastatic lesion (green triangles). See Supplementary Fig.2D-E for representative images.

In the HEp3 model we showed that macroH2A1 levels were expressed at higher levels in solitary DCCs and small clusters (**Fig.1**). We next tested whether solitary DCCs present in the LN of HNSCC patients also show increased levels of macroH2A compared to PT samples. While LN samples are not commonly processed for the detection of solitary DCCs, we were able to gain access to a small repository of samples where clinically negative. LN FFPE samples were collected during HNSCC primary tumor removal surgeries and confirmed to contain solitary cytokeratin positive DCCs^39, 40^. IF staining revealed that both macroH2A1 and macroH2A2 signals were easily detectable in LN DCCs (**Fig.2C-D**), and at least 2-to 5-fold higher in protein expression in solitary DCCs compared to non-paired PT (**Fig.2E-F**). When normalizing for abundance in PTs across the patient cohort 1 (US-based patients) and cohort 2 (Germany-based patients) taking the PT in both cohorts as baseline expression (see Methods), macroH2A signals were also upregulated in solitary DCCs *vs*. LN metastasis **(Fig.2E-F)**. Despite being a small sample set and using normalization to account for inter-cohort differences, this analysis suggests that solitary or small clusters of DCCs show increased macroH2A levels compared to PT and LN metastatic samples.

We also explored whether macroH2A1 was differentially expressed in DCCs isolated from the bone marrow of prostate cancer patients with no evidence of disease (NED) for up to 18 years or with advanced metastatic disease (ADV) that we previously analyzed for other dormancy markers^7, 41^. Interestingly just like for TGFβ2 and NR2F1^7^, we found that macroH2A1, but not macroH2A2, mRNA levels were significantly higher in DCCs from the NED vs the ADV group of patients (**Supplementary Fig.2E)**. While in small patient sample sized cohorts, these data support that macroH2A variant expression may be a feature of dormant disease across different human cancers.

### MacroH2A overexpression induces a quiescent phenotype in malignant HNSCC cells

We next tested whether macroH2A1 is essential for the cellular dormancy in HEp3 variants. We applied CRIPSR-Cas9 knockout or shRNA methods to deplete macroH2A1 in spontaneously dormant D-HEp3 cells (macroH2A1 is the only variant expressed in D-HEp3 and T-HEp3 cells) (**Supplementary Fig.3A-C**), and inoculated the engineered D-HEp3 cells *in vivo* on CAM for 7-day incubation, where they enter a long lasting G0/G1 arrest that can last for at least two months^29,3, 42^. These experiments revealed that the dormant phenotype of D-HEp3 cells was not interrupted by macroH2A1 knockout or knockdown as tumor nodules remained small (20-50 mm^3^) compared to proliferative T-HEp3 cells that can reach >10 times that volume at the same time^29^ (**Fig.3A**). This suggests that macroH2A1 is not essential for the dormant phenotype of D-HEp3 cells. However, solitary DCCs in LNs upregulated macroH2A variants (**Fig.2E-F**).

**Figure 3.**
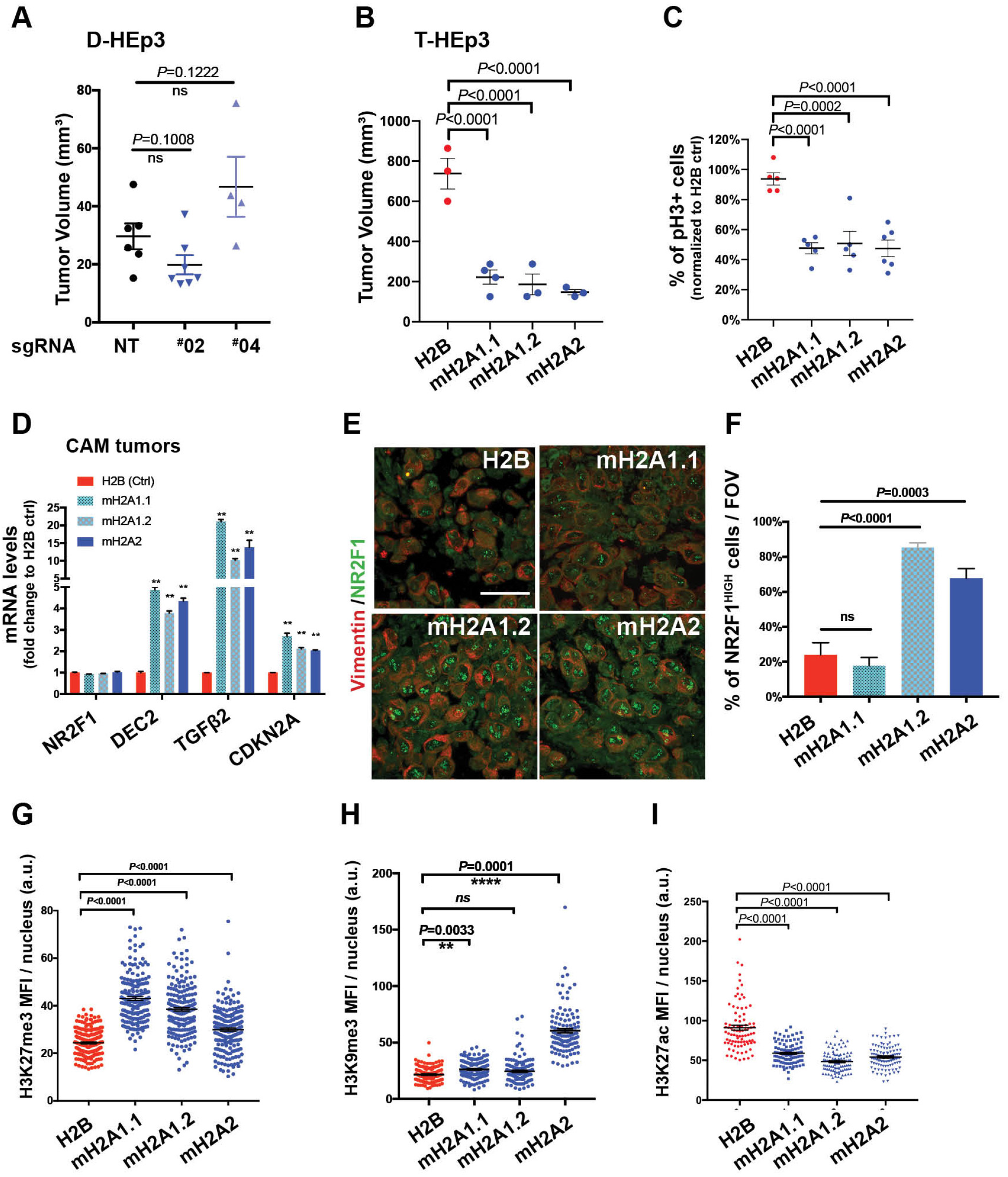
MacroH2A overexpression induces a quiescent phenotype in malignant HNSCC cells. (A) Tumor volume measurement of D-HEp3 CAM tumors with non-targeting guide RNA (NT) and 2 different guide RNAs targeting macroH2A1 (#02, #04). D-HEp3 cells were inoculated onto CAM (5×10^5^ cells per animal) for 7-day incubation. (n =6 (NT), 7 (#02), 4 (#04) tumors). Unpaired Student’s t-test. (B) Tumor volume measurement of T-HEp3 CAM tumors with H2B-GFP (Ctrl) and different variants of macroH2As construct expression. T-HEp3 cells were inoculated onto CAM (1.5×10^5^ cells per animal) and transplanted into a new chicken embryo CAM for the 2^nd^ week (n =3 (H2B), 4 (mH2A1.1), 3 (mH2A1.2), 3 (mH2A2) tumors). (C) Quantification of percentage of H3S10phos positive cells on CAM tumors, normalized to that observed in H2B control tumors. 200∼300 cells were assessed per tumor section; 5 tumors were assessed per condition. Ordinary one-way ANOVA test. (D) qPCR analysis of dormancy gene mRNA levels in the indicated tumors grown on CAM for 1 week. PCR in triplicate, mean + s.e.m. Multiple t tests with false discovery rate approach. (E-F) Images and quantification of the indicated T-HEp3 CAM tumor sections stained for NR2F1 (green). Tumor cells were also stained with human vimentin (red). Scale bar, 50um. (n>100 cells assessed per section, 3 sections assessed from 3 different tumors per condition), ordinary one-way ANOVA test. (G-I) Quantification of MFI from repressive (H3K27me3, H3K9me3) and active (H3K27ac) histone modification IF in the indicated T-HEp3 CAM tumor sections, n ≈200 cells assessed from 3 different tumors per condition.

Thus, we tested whether inducible overexpression of macroH2A might be sufficient to initiate a cellular dormancy mechanism. To test this possibility, we generated T-HEp3 cells overexpressing each individual isoform of macroH2A or the histone H2B as control (as GFP fusions; **Supplementary Fig.3D**) and subsequently inoculated the cells *in vivo* on the CAM and followed serial passages of tumors for two weeks, which is sufficient to confirm a dormancy-like phenotype^6, 7^. Strikingly, overexpression of all macroH2A variants was sufficient to suppress tumor growth *in vivo* (**Fig.3B**). This phenotype was due to reduced proliferation rather than apoptosis, supported by significantly reduced H3S10 phosphorylation and no change in cleaved caspase-3 abundance in macroH2A overexpressing cells (**Fig.3C, Supplementary Fig.3E-F**).

MacroH2A-induced growth arrest was associated with upregulation of TGFβ2 and DEC2, as well as the CDK inhibitor, CDKN2A/p16 (**Fig.3D**). While TGFβ2 can induce macroH2A1 expression in T-HEp3 cells (**Fig.1E**), macroH2A variants also induced TGFβ2, suggesting that these genes may be part of an autocrine feedforward loop that initiates and maintains dormancy reprogramming of tumorigenic cancer cells. In contrast, the transcript level of NR2F1, was not affected by ectopic macroH2A variant expression. However, IF analysis revealed that macroH2A overexpression triggered a robust accumulation of NR2F1 signal in the nucleus (**Fig.3E-F**), where it is commonly found in D-HEp3 cells^7^(**Supplementary Fig.3G)**, in spontaneously dormant DCCs in mice and humans^7, 15^ or cells treated with an NR2F1 agonist^32^. This change was significant when macroH2A1.2 or macroH2A2, but not macroH2A1.1 was overexpressed (**Fig.3E-F**), suggesting different mechanisms driven by the variants in this particular phenotype. NR2F1 and macroH2A may be part of a self-reinforcing mechanisms since an NR2F1 agonist can also induce macroH2A1 expression (**Fig.1F-I**).

Previous studies demonstrated that dormant tumor cells with NR2F1^high^ signal display a repressive chromatin state, with increased H3K9me3 and H3K27me3 nuclear signal^7^. Accordingly, we detected an enrichment of repressive chromatin modifications by overexpressing macroH2A variants. These included an increased H3K27me3 nuclear signal and decreased H3K27ac nuclear signal when overexpressing all individual macroH2A variants; macroH2A1.1 overexpression significantly increases the H3K9me3 nuclear signal, however macroH2A2 overexpression is much more robust for H3K9me3 (**Fig.3G-I, Supplementary Fig.3H-J**). These data support that macroH2A expression in malignant cancer cells can induce a growth arrest program displaying hallmarks of a repressive chromatin reorganization like spontaneous dormant tumor cells.

### MacroH2A2-induced dormancy displays features of spontaneous dormancy and senescence

While macroH2A2 expression is undetectable in both T-HEp3 cells and D-HEp3 cells at the mRNA and protein levels (**Supplementary Fig.1B-C**), its overexpression was sufficient to induce a dormancy-like program. These data led us to focus our attention on this macroH2A variant, as its downregulation is also associated with melanoma metastasis in patients, and acts as the strongest barrier to somatic cell reprogramming of all three macroH2A isoforms^22, 28^. To decipher how macroH2A2 is driving cells into a dormancy-like state, we tested whether NR2F1 or DEC2 were required for macroH2A2-induced growth arrest *in vivo*. Knockdown of either TF in macroH2A2 overexpressing cells did not reverse macroH2A2-induced growth arrest, indicating that while induced, these TFs were not necessary for macroH2A2-induced dormancy (**Supplementary Fig.4A-D**).

This suggested that macroH2A2 may activate a unique gene expression program when inducing dormancy. Transcriptomic profiling by RNA sequencing (RNA-seq) of T-HEp3 cells either overexpressing macroH2A2 or H2B as control identified 914 differentially expressed genes (DEGs), when using a log2 fold change cut-off of 0.5 (**Fig.4A, Supplementary Fig.4E-F, Supplementary Table 1**). Pathway’s analysis using GSEA (**Supplementary Tables 2-3**) or Enrichr, a comprehensive resource for curated gene sets dataset^43^ (**Supplementary Table 4**) supported the tumor growth arrest phenotype. GSEA analysis revealed E2F targets and MYC targets as top negatively regulated hallmark gene sets (**Fig.4B-C**). In addition, DNA replication, DNA repair, DNA strand elongation and other S phase-associated terms were enriched with downregulated gene set *via* Enrichr (**Supplementary Table 4**), consistent with the potent growth arrest *in vivo*. Enrichr analysis with Molecular Signatures Database (MSigDB) Hallmark Gene Set also revealed that genes commonly associated with fatty acid metabolism were suppressed by macroH2A2 overexpression. Our global DEG analysis supports that lack of macroH2A2 in malignant cells allows for active proliferation and potentially activation of lipid metabolism, which may be linked to metabolic reprogramming that happens in the transition from quiescence to proliferation^44^. In contrast, the pathway enrichment analysis of up-regulated genes (n=485) showed epithelial-to-mesenchymal transition (EMT), inflammatory response and circadian rhythm regulation (**Supplementary Table 4**). The exact role of the circadian rhythm gene regulation is unclear but DEC2/BHLHE41 is an important clock gene^45^, and loss of circadian regulation can result for example in MYC-driven tumors due to loss of growth suppression^46^.

**Figure 4.**
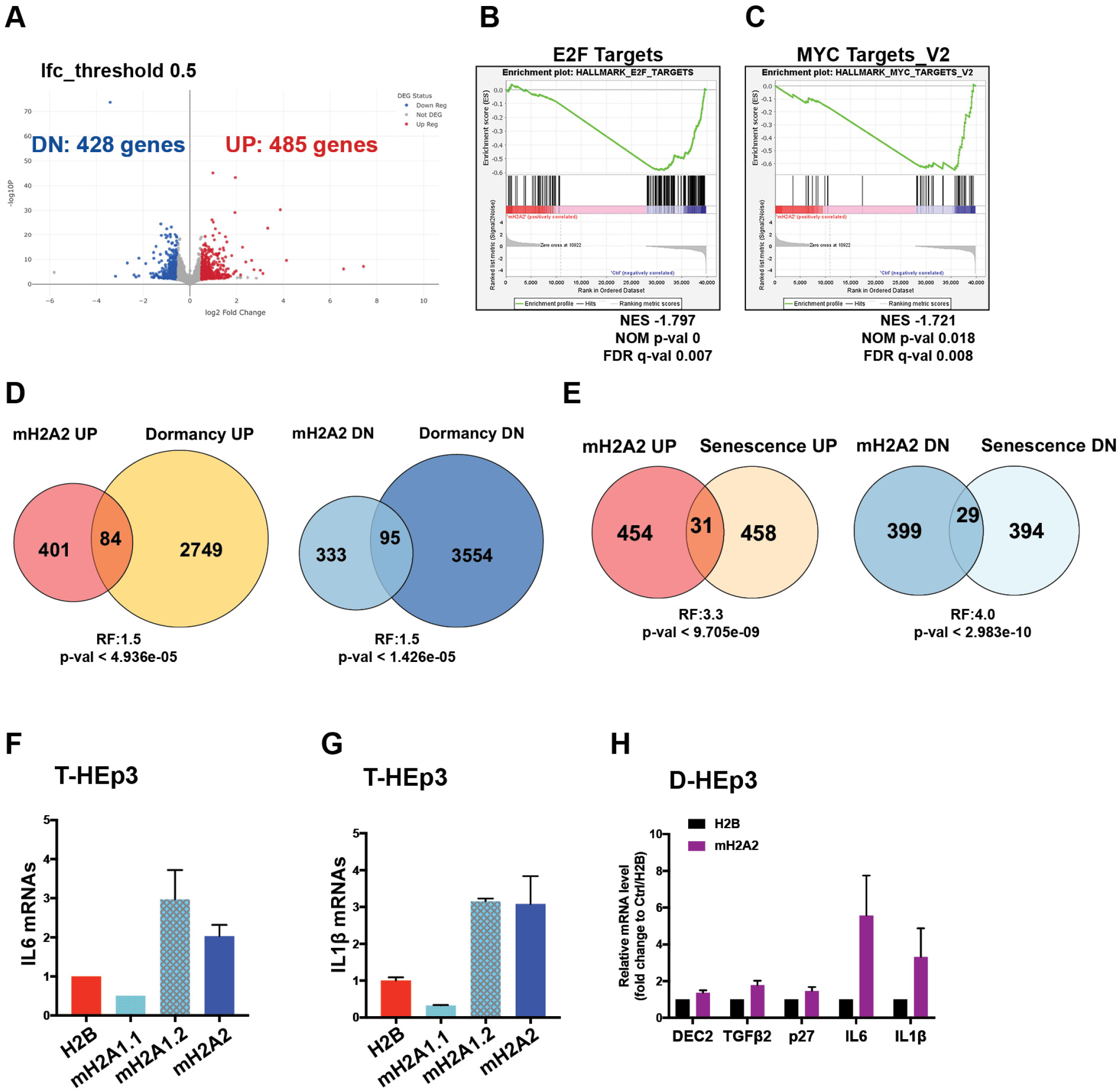
MacroH2A2-induced dormancy displays features of spontaneous dormancy and senescence. (A) Volcano plot displays the differentially expressed genes (DEGs) detected by RNA-seq from T-HEp3 cells with mH2A2 vs. H2B overexpression. Log2foldchange cut-off is 0.5; 486 up-regulated genes, 428 down-regulated genes. (B-C) Top negatively correlated Hallmark Gene Sets using RNA-seq data via Gene Set Enrichment Analysis. (D) Venn diagrams show the comparison of upregulated and downregulated genes in D-HEp3 with up and downregulated genes in macroH2A2 overexpressing cells. Representation factor:1.5, *P*<4.936e-05, *P*<1.426e-05 (E) Venn diagrams show the comparison of up- and down-regulated genes in oncogene-induced senescent fibroblasts (IMR90) with up- and down-regulated genes in macroH2A2 overexpressing T-HEp3 cells. Representation factor: 3.3, *P*<9.705e-09; Representation factor: 4.0, *P*<2.983e-10. (F-G) qPCR analysis of IL6 and IL1β transcript levels in T-HEp3 cells overexpressing different macroH2A variants. (H) qPCR analysis of dormancy and senescence gene mRNA levels in the indicated D-HEp3 cells with mH2A2 overexpression or H2B as control. PCR in triplicate, mean + s.e.m.

Enrichr analysis with MSigDB Hallmark Gene Set also (**Supplementary Table 4**) revealed that genes commonly associated with inflammatory response or TNF-alpha signaling via NF-kB programs are up-regulated by macroH2A2 overexpression. A similar trend was detected when using GSEA analysis (**Supplementary Table 2**). This suggested that potentially, cytokines, which are commonly associated with inflammation, might be upregulated by macroH2A2. Interestingly some of these cytokines such as IL6 and IL1β have been linked to the senescence associated secretory phenotype (SASP)^47^. These associations and the fact that macroH2A is enriched in SAHF^16^, led us to compare the macroH2A2 gene expression profile to those of dormant D-HEp3 vs. isogenic proliferative T-HEp3 cells (GSE172115) and IMR90 fibroblasts undergoing oncogene induced senescence vs. proliferative fibroblasts^48^ **(Fig.4D-E)**. These comparisons revealed that only a small fraction of genes regulated in dormant D-HEp3 cells is also regulated by macroH2A2 (**Fig.4D and Supplementary Table 5**). When we performed the same analysis comparing with senescent cells, we found that macroH2A2 overexpression resulted in an overlapping program, but still limited set of genes (**Fig.4E and Supplementary Table 6**). GSEA analysis of the macroH2A2 regulated genes compared to a senescence and autophagy signature (**Supplementary Table 3**), although not statistically significant, also showed a partial overlap with this senescence signature. This argues that macroH2A2 is only regulating a sub-group of senescence regulated genes, explaining why GSEA analysis is not significant. Nonetheless, macroH2A2 induced a strong proliferative arrest, suggesting that a combination of dormancy and senescence sub-programs regulated by this unique histone variant may be sufficient to induce a long-term arrest in malignant cells.

When considering the common genes upregulated by macroH2A2 and senescent cells, which also included genes enriched in the Senescence and Autophagy in cancer pathway (**Supplementary Table 3**), we found that several cytokines induced by macroH2A2 and found in the SASP, or other secreted molecules also associated with senescence and aging, were not upregulated in dormant D-HEp3 vs. T-HEp3 cells (**Supplementary Tables 5-6**). These included IL1β, IL6, IL11, GDF15, GDNF, TNC, LAMA4. We found that overexpression of macroH2A1.2 and macroH2A2, but not macroH2A1.1, increased IL6 and IL1β transcript levels via qPCR **(Fig.4F-G**). These data suggest that macroH2A2 may be inducing a specific subset of senescence-associated genes (IL1β, IL6). Overexpression of macroH2A2 in dormant D-HEp3 cells, in addition to reinforcing the expression of DEC2, TGFβ2 and p27, induced >5-fold induction in IL1β and IL6 **(Fig.4H)**, suggesting that even in cells carrying a quiescence program of dormancy, a gain of macroH2A2 could induce senescence-associated genes. We conclude that macroH2A2 induces a dormancy-like program with components of quiescence and senescence gene programs in malignant T-HEp3 cells.

Since primary lesions can still express macroH2A variants (**Fig.2**), we wondered whether the expression levels of macroH2A and the enrichment of the macroH2A-regulated gene set described above would inform on overall and relapse free survival in HNSCC patients^49^. Analysis of these parameters revealed that over a four-year follow-up, patients carrying primary tumors with a high macroH2A2 gene signature (see Supplementary Fig.4 legend and Supplementary Table 1 for details on the genes used to define the signature) displayed longer overall survival and relapse free periods, albeit the latter was not powered for significance (**Supplementary Fig.4G-H**). These data suggest that tumors or cells within a tumor carrying a high macroH2A2 expression level may spawn DCCs with a higher propensity to enter dormancy.

### MacroH2A2 promotes the accumulation of long-lived dormant DCCs restricting their ability to initiate proliferative metastasis

HEp3 cells can form overt metastases in lungs and lymph nodes in either experimental or spontaneous metastasis models^6, 36^. We next studied whether macroH2A2 could influence DCC fate *in vivo*. First, we performed an experimental metastasis assay with T-HEp3 cells expressing either H2B or macroH2A2 GFP fusions. Three weeks after tail-vein injection in nude mice, the lungs were harvested and processed for IF analysis. Using vimentin antibody^6, 7^ that preferentially detects human cells, we scored the frequency of solitary DCCs (including single cells and less than 8-cell clusters), micro-metastasis and large metastatic events in each animal (**Fig.5A**). MacroH2A2 overexpression led to a higher frequency of solitary DCCs and fewer micro- and macro-metastases in the lungs, compared to the control group (**Fig.5B**). DCCs overexpressing macroH2A2 (confirmed by GFP expression) displayed a higher proportion of NR2F1-positive cells (also observed in **Fig.3E**), which was inversely correlated with the frequency of H3S10phos-positive DCCs, compared to the H2B-GFP DCCs (**Fig.5C-E**). Thus, macroH2A2 can induce DCC growth arrest associated with the dormancy marker NR2F1. These data argue that macroH2A2 promotes the accumulation of dormant DCCs at the expense of proliferative metastases.

**Figure 5.**
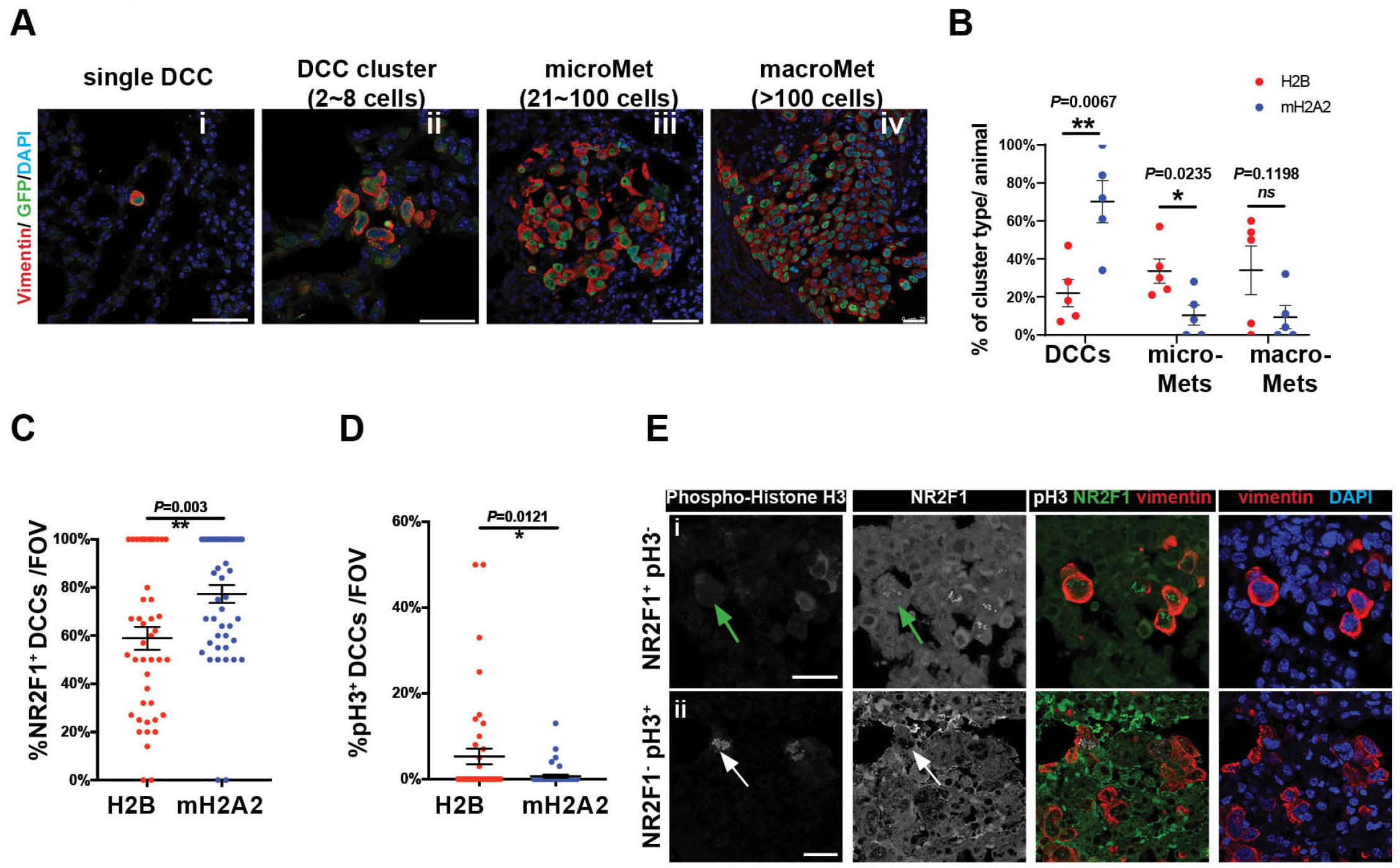
MacroH2A2 overexpression promotes the accumulation of quiescent DCCs to prevent metastatic growth. (A) Experimental metastasis assay. T-HEp3 cells with GFP-mH2A2 or GFP-H2B expression were tail vein injected in nude mice. Three weeks later, the lungs were retrieved and processed for FFPE sections. n = 5 animals were accessed per condition. Representative IF images of single DCC, DCC cluster (< 8 cells), micro-Met and macro-Met were co-stained with GFP, human vimentin (red) and DAPI (blue). Scale bar, 50um. (B) Quantification of DCC, micro-Met and large-Met event frequency in the freshly resected lungs in each animal. Singles cells and small clusters (<8 cells) are all included as DCC events. Red dots annotate control animals with H2B expression, while blue dots annotate experimental animals with mH2A2 overexpression. Mean ± s.e.m. Two-tailed Unpaired t-test. (C-D) Quantification of IF staining for NR2F1, H3S10phos and human vimentin in DCCs found in the lungs per condition. *n=* 80 fields of view pooled from 5 animals per condition. (E) Representative IF images of NR2F1 (green), human vimentin (red) and H3S10phos (grey) in lung DCCs. Green Arrow, NR2F1 positive cell; White Arrow, pH3 positive cell. Scale bar, 25 um.

To further test the dormancy-inducing function of macroH2A2 in DCCs, we next performed a spontaneous metastasis assay (**Fig.6A**) using engineered Tet-ON-inducible macroH2A2-GFP T-HEp3 cells (**Supplementary Fig.5A-B**). This assay allows the examination of whether an adjuvant “therapeutic” induction of macroH2A2, as it has been tested for oncogene de-induction in inducible mouse models^50, 51^, could induce and/or maintain dormancy after cancer cells have completed all steps of dissemination with no macroH2A2 expression. To this end, 5×10^5^ T-HEp3 Tet-ON-macroH2A2-GFP cells (per animal) were injected subcutaneously into nude mice and when primary tumors reached ∼500mm^3^ they were surgically removed. Animals were then provided with oral treatment (“therapeutic” phase) of doxycycline (2mg/mL)^52^. Six weeks after surgery, we found that ∼57% of animals in the no DOX treatment control group developed local recurrences in the surgery margins where dormant residual cells persist and fuel recurrences^7^. In contrast, animals with inducible expression of macroH2A2 displayed greatly reduced local recurrences, with only 11% of animals relapsing (**Fig.6B-C**). Further, the local recurring lesion expressing macroH2A2 grew at a much slower rate than those developed in the negative control group (**Fig.6B-C**). This indicates that macroH2A2 prevents local residual tumor cells in surgery margins of the primary tumor from reactivating.

**Figure 6.**
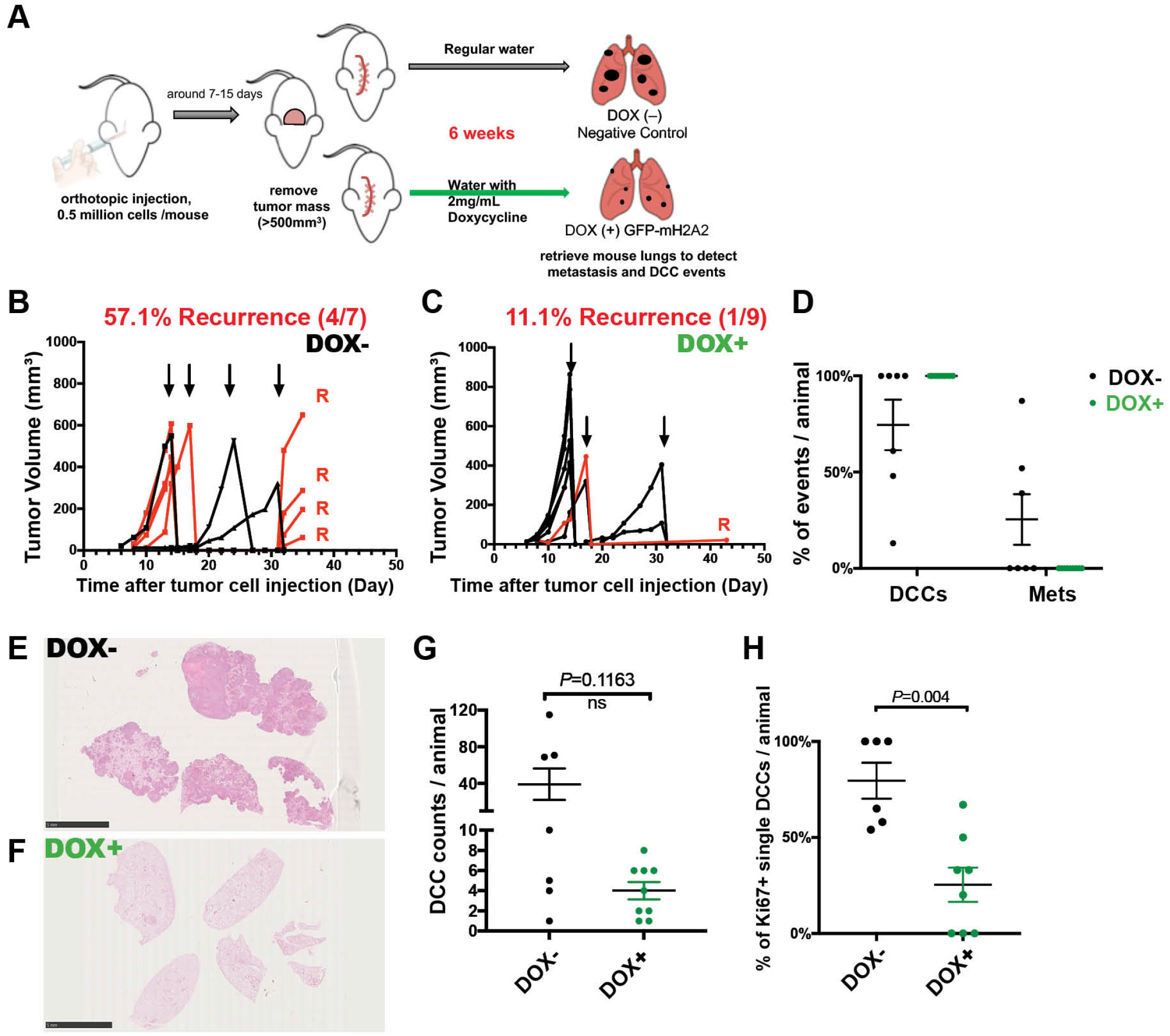
Induction of macroH2A2 after tumor cell dissemination restricts their ability to initiate proliferative metastasis. (A) Scheme of spontaneous metastasis assay. Tet-ON-inducible macroH2A2-GFP T-HEp3 cells were s.c. injected in nude mice. Tumors were surgically removed when the size reached ∼500 mm^3^ then the mice were provided with oral treatment (2 mg mL^-1^ DOX water, replenished every 48hr) to induce the expression of macroH2A2-GFP or regular water without DOX. Six weeks later, the lungs were retrieved and processed for FFPE sections. (B-C) Measurement of the primary tumor volumes after the orthotopic injection. Arrows indicate the primary tumor removal times. Red lines annotate animals with local tumor recurrence after the surgery. (D) Frequency measurement of lung DCC and metastasis events in each animal. Single cells and doublets are both considered as DCC events. Black dots represent animals without doxycycline treatment (DOX^-^) (n=7), while green dots represent animals with oral doxycycline (DOX^+^) (n=9). (E-F) Representative H&E images of animal lungs with metastasis growth in DOX^-^ group and those with no evidence of metastasis in DOX^+^ group. Scale bar, 5mm. (G) Counts of DCC events in each animal, 12 tissue sections pooled from each animal. (H) Quantification of IF staining for Ki67-positive solitary DCCs found in the lungs. Mann-Whitney test.

At the same end point, we additionally quantified the frequency of lung DCC and metastasis in animals with or without DOX treatment using IF with vimentin and Hematoxylin and Eosin staining (**Fig.6D-F**). We found that inducible expression of macroH2A2 suppressed overt metastatic growth (**Fig.6D-F**) that was present in 3 out of 7 control group animals (**Supplementary Fig.5C-D**). This was evidenced by inducible expression of macroH2A2 resulting in lungs only carrying solitary and small clusters of DCCs, with not even micro-metastases (20-100 cells/lesion) being detectable (**Fig.6D**). No statistically significant difference in the number of solitary DCCs detected in two groups was detected, suggesting that seeding was similar. However, the proportion of proliferative DCCs (Ki67 positive) is significantly higher in the negative control group (**Fig. 6G-H**), arguing that macroH2A2 holds DCCs in a dormant state associated with a NR2F1^hi^ /p-H3^lo^/Ki67^lo^ profile. Our results indicate that restored macroH2A2 expression is sufficient to keep DCCs in the state of dormancy in lungs.

## Discussion

Clinical dormancy can be a long-lived event lasting many years^2^. This led us to hypothesize that if DCCs persist in a long-term growth arrest with perhaps punctual divisions, like quiescent adult stem cells^53^, convergence of both microenvironmental and epigenetic mechanisms might be necessary to maintain this phenotype. Previous work from our labs suggested that mechanisms of growth suppression, dormancy induction and metastasis suppression^7, 28, 54, 55^ are associated with transcriptional and chromatin remodeling events. In particular, a repressive chromatin state was found in HNSCC dormant cancer cells linked to the TF NR2F1^7^, and in melanoma cancer cells, metastasis-initiating capacity was suppressed by overexpression of macroH2A variants^28^. However, whether metastasis suppression by macroH2A variants and dormancy induction were functionally linked was unknown.

Our analysis of the above question revealed that TGFβ2, a pro-dormancy microenvironmental cue found in the ECM and in soluble form^6, 31, 36^ was able to induce macroH2A1 expression and a dormant phenotype. The p38α/β pathway, which also responds to other microenvironmental signals such as TGFβ1/2^6, 31^, BMP7^8^ and retinoic acid^7, 56^, also induces and/or maintains expression of macroH2A1. Activation *via* an agonist of the orphan nuclear receptor NR2F1, also regulated by BMP7 and retinoic acid^7^, again resulted in macroH2A1 variant upregulation. Thus, it is possible that niche-derived cues that control growth arrest programs of quiescence or differentiation can upregulate macroH2A1 to induce cancer cell growth suppression. However, in the D-HEp3 dormancy model^29^, macroH2A deficiency was insufficient to reverse dormancy. We reasoned that other transcriptional and signaling nodes active in these cells and known to induce dormancy^7, 14^ still exert a growth suppressive effect even in the absence of macroH2A. Nevertheless, stable or inducible overexpression of macroH2A1 or macroH2A2 in malignant HEp3 cells was sufficient to induce a dormancy-like program. Interestingly, macroH2A variant upregulation strongly upregulated TGFβ2, DEC2 and p16, but this induced dormancy program did not depend on the known dormancy TFs, DEC2 and NR2F1 for reasons that remain unknown. Of note, NR2F1 was not induced at the mRNA level, but accumulated robustly in the nucleus upon macroH2A variant expression, supporting that it might recruit NR2F1, and possibly other nuclear factors to enforce the growth arrest program.

The above results argued that an additional mechanism was operating downstream of macroH2A to induce growth suppression. RNA-seq analysis revealed that the growth arrest induced by macroH2A2 only partially overlapped with quiescence and senescence gene expression programs found in dormant D-HEp3 cells^7, 57^ or senescent fibroblasts^48^, respectively. Some of the gene expression programs were indeed expected, such as repression of MYC- and RAS-regulated programs. While macroH2A2-induced only a subset of quiescence- and senescence-associated genes, their combination was associated with a strong suppression of metastasis and local recurrence initiation by holding solitary DCCs in a dormant-like state. This program involves cytokines uniquely regulated by macroH2A2 such as IL1β, IL6, IL11, GDF15 and GDNF. Although unknown, we hypothesize that these may cooperate with TGFβ2 signaling to induce dormancy. The transcriptional regulators executing these programs downstream of macroH2A2 remain to be identified.

A remarkable finding was that overexpression of macroH2A1.2 and macroH2A2, but not macroH2A1.1, was sufficient to induce the expression of mRNAs commonly found to be produced in senescence (e.g., IL1β, IL6). Interestingly, only the macroH2A1.1 macro domain is described to bind ADP-ribose metabolites and ADP-ribosylated proteins, which is not present in the other variants^58, 59^ and may have a transcriptionally-independent metabolism regulating function^60^. Thus, the selective function of macroH2A1.2 and macroH2A2 found in this model may be attributed to transcriptional regulation functions of these two variants. For example, our data suggests that only macroH2A1.2 and macroH2A2 induce NR2F1 nuclear accumulation. We believe, however, that the induction of SASP genes is likely an indirect effect of the macroH2A-induced growth arrest, rather than direct macroH2A binding at these loci and inducing their expression. In fact, macroH2A1 has been shown to be specifically removed from SASP gene chromatin during OIS^61^.

The literature has implicated loss of macroH2A2 as a common event in cancer^25, 27, 28^, further supporting a restrictive role for cancer progression and metastasis initiation in our studies. Of note, however, the contribution of each variant to tumorigenesis appears highly cell-type and context dependent^62^, and an intriguing finding along these lines is that while the T-HEp3 cells express some macroH2A1, they were highly proliferative and metastatic at baseline. Nevertheless, overexpression of the same variants was able to reprogram the cells into growth arrest. This suggests that malignant cells may restrict the access of macroH2A variants to loci that control growth suppression programs but that these loci can be re-engaged by available macroH2A molecules. Further exploration of these mechanisms may allow “repositioning” macroH2A in malignant cells with basal level expression to activate these quiescence and senescence partial programs.

The fact that macroH2A histone variant expression can persist to a certain level without causing growth suppression was also observed as both macroH2A1 and macroH2A2 are present in aggressively proliferative lesions in head and neck primary tumor patient samples. However, an important distinction was that the solitary and potentially dormant cells robustly upregulate the levels of macroH2A variants, and the expression of all variants significantly decreases in proliferative metastatic lesions. These data suggest that during progression, cancer cells can spontaneously upregulate macroH2A variants to adapt to the switch from proliferation to quiescence, but that metastatic growth is associated with a downregulation of all variants consistent with our findings in melanoma^28^.

Our results reveal that the macroH2A histone variants play a significant role in the mechanisms of metastasis suppression. We reveal that these mechanisms suppress metastasis *via* dormancy induction but with unique and a limited set of genes found in senescent fibroblasts but not spontaneously dormant D-HEp3 cells. These results suggest that in order to induce and maintain dormancy, activating a subset of diverse components of distinct growth arrest programs may be sufficient. In this study, we reveal that macroH2A1/2 variants function could be exploited for such a purpose and that their detection (or a macroH2A-regulated signature) in primary lesions or in DCCs might be useful biomarkers of patients with better prognosis and/or dormant DCCs, and could be used in clinical trials to phenotypically profile DCCs^31, 56^. Given that senescent cells can also be cleared by innate immune cells^63^ our results open the possibility the macroH2A2 induced DCC dormancy could eventually lead to eradication of these residual cells *via* the immune system. Future studies of macroH2A manipulation in DCCs, and their interaction with the tumor microenvironment, will reveal such answers.

## ACKNOWLEDGEMENTS

We thank the Aguirre-Ghiso, Sosa and Bernstein labs for helpful discussions. This work was supported by grants from National Institutes of Health (NIH)/National Cancer Institute (NCI) (CA109182, CA216248, CA218024, CA196521 to JAAG and R01CA154683, CA218024 to EB). JAAG is Samuel Waxman Cancer Research Foundation Investigator. RNA-seq was supported in part through the Oncological Sciences Sequencing Core supported by Tisch Cancer Institute (TCI) of the Icahn School of Medicine at Mount Sinai (ISMMS) Cancer Center Support Grant P30 (CA196521), Scientific Computing supported by the Office of Research Infrastructure of the NIH under award number S10OD026880 to ISMMS, and the Mount Sinai Genomics Technology Facility, and Deniz Demircioglu from the Bioinformatics for Next Generation Sequencing (BiNGS) shared resource facility within the TCI at ISMMS, partially supported by P30 (CA196521).

## AUTHOR CONTRIBUTIONS

DS designed, planned, and conducted experiments, analyzed data, and wrote the manuscript; DS and DF performed *in vivo* mouse experiments and DS further processed *in vivo* material; DS, SC, DH and DKS performed RNAseq experiments and analyzed data; BM, WW, and CS provided human tissue samples and assisted tissue anatomical analysis. JAAG conceived the project, designed experiments, analyzed data, and wrote the manuscript. JAAG and EB analyzed data, secured funding, and wrote and edited the manuscript.

## FIGURE LEGENDS

**Supplementary Fig.1 MacroH2A differential expression in HNSCC PDX cell lines**

(A) Diagram depicts the growth phenotype of T-HEp3 and D-HEp3 cell variants on CAM system. (B)Snapshots of the UCSC genome browser (hg19) showing RNA-seq, H3K27ac and H3K27me3 signal at the MACROH2A1 (upper) and MACROH2A2 (lower) loci in T-HEp3 and D-HEp3, respectively. (y axis = reads per kilobase per million reads) Bar graph (right) indicates expression values (Median-ratios normalized reads) in corresponding cells. (C) Western blots show mH2A2 expression levels in different head and neck squamous cell carcinoma cell lines. Histone H4 is used as housekeeping gene loading control. mH2A2 molecular weight is approximately 40k Da. (D) Western blots show decreased expression levels of phosphoATF2 and p27, due to the inhibition of p38 α/β activity by SB 203580 (5uM, 48hrs) treatment in serum free medium *in vitro*. (E) Cell number count per tumor of T-HEp3 CAM tumors with or without 10ng/mL TGFβ2 treatment for 6 days. 2 ×10^5^ cells were inoculated per CAM. (F) The scheme of the experimental metastatic assay shown in Figure 1J-L.

**Supplementary Fig.2 Heterogeneous expression of macroH2As in HNSCC and prostate cancer patients**

(A-B) Box plots show the frequency of macroH2A1^HIGH^ (A) or macroH2A2^HIGH^ (B) cells in paired MS-PT and MS-LN Met patient samples. *P* value, Mann-Whitney test. (C) Image analysis in details and representative images from one patient ^#^58305. FFPE samples were stained with pan cytokeratin (Pan-CK) in red, macroH2A1 in green and Histone H3 in blue. *n*=6 fields of view were assessed in each condition. Mean fluorescent intensity of macroH2A1(green) or Histone H3 (blue) was quantitatively measured in ImageJ. Cells with macroH2A1/H3 intensity ratio higher than 0.75 were considered as macroH2A1^HIGH^. (D) Image analysis in details and representative images from one patient ^#^59152. FFPE samples were stained with pan cytokeratin (Pan-CK) in red, macroH2A2 in green and Histone H3 in blue. n=6 fields of view were assessed in each condition. Mean fluorescent intensity of macroH2A2 (green) or Histone H3 (blue) was quantitatively measured in ImageJ. Cells with macroH2A2/H3 intensity ratio higher than 0.6 were considered as macroH2A2 ^HIGH^. Scale bar, 50um. (E) Relative mRNA levels of macroH2A1 and macroH2A2 in isolated bone marrow DCCs from prostate cancer patients with no evidence of disease (NED) and with metastatic disease or biomedical recurrence following primary treatment (advanced disease, ADV).

**Supplementary Fig.3 MacroH2A overexpression induces tumor growth arrest with repressive chromatin reorganization**

(A) Western blots show the CRISPR-Cas9 knockout of macroH2A1 in D-HEp3 cells. (B) Western blots show partial knockdown of macroH2A1 via shRNAs in D-HEp3 cells. (C) Cell number counts of individual D-HEp3 CAM tumors with scramble shRNA (NC) and 2 different shRNAs to macroH2A1 (^#^90, ^#^93). D-HEp3 cells were inoculated onto CAM (5×10^5^ cells per animal) for one week incubation. (n =4 (NC), 4 (^#^90), 3 (^#^93) tumors). Two-tailed Unpaired t-test. (D) Western blots show the expression of GFP fused different macroH2A variants in T-HEp3 cells, comparable to the endogenous level of indicated proteins. (E-F) Images and quantification of the indicated T-HEp3 CAM tumor sections stained for cleaved Caspase 3 (green). Tumor cells were also stained with human vimentin (red) and DAPI (blue). Scale bar, 50um. (G) Representative IF image of D-HEp3 CAM tumor section stained for NR2F1 (green). Tumor cells were also stained with human vimentin (red). (H-J) Representative IF images of the indicated T-HEp3 CAM tumor sections stained for repressive or active histone marks-H3K27me3(H), H3K9me3(I), H3K27ac(J) in green and vimentin in red. Scale bar, 50um or 25um.

**Supplementary Fig.4 MacroH2A2-induced dormancy is independent of NR2F1 and DEC2, but its gene expression signature correlated with long-term dormancy in patients**

(A) qPCR analysis of NR2F1 transcript level after 48hr treatment with scramble RNA or two separate siRNAs. Two-tailed Unpaired t-test. (B) T-HEp3 Tumor growth on CAMs for 5 days. T-HEp3 cells were treated with NR2F1 knockdown (siRNA NR2F1#4) for 48hrs before CAM inoculation. (C) qPCR analysis of DEC2 transcript level after 48hr treatment with scramble RNA or 2 separate siRNAs to DEC2. Two-tailed Unpaired t-test. (D) Tumor growth on CAMs for 5 days of T-HEp3 cells with DEC2 knockdown (siRNA DEC2#2). (E) PCA plot generated with RNAseq analysis from T-HEp3 cells with mH2A2 overexpressing vs. H2B expressing. (F) Heatmap of DEGs detected by RNAseq. 486 up-regulated genes, 428 down-regulated genes; adj.*p* value<0.05, Log2 FC>0.5 or< -0.5. (G-H) Overall survival (G) and relapse free survival (H) were analyzed using RNAseq data of KM plotter in HNSCC patients with a 4-year follow-up. The patients are stratified in two groups (high expression in red and low expression in black) according to the expression profiles of mH2A2-induced top 52 up-regulated genes. P-values for the significance of the difference between high and low expression were calculated using the log-rank test.

**Supplementary Fig.5 MacroH2A2 restricts proliferative metastasis**

(A) Western blots show the Tet-ON inducible macroH2A2-GFP expression is induced via different doses of doxycycline after 48 hours treatment *in vitro* cell culture. (B) Tumor volume measurement of individual T-HEp3 CAM tumors with or without inducible GFP-mH2A2 expression after 6-day incubation on CAM. 2.0 ×10^5^ cells T-HEp3 cells were inoculated onto CAM. (C-D) Hematoxylin and eosin stain of animal lungs retrieved 6 weeks after primary tumor removal. Scale bar, 5mm. R, indicates animals with local recurrences developed after primary tumor removal.

### Supplementary Tables

**Supplementary Table1**. Differentially expressed genes (DEGs) in T-HEp3 cells overexpressing mH2A2 vs. H2B.

**Supplementary Table2**. GSEA analysis of hallmark pathways in T-HEp3 cells overexpressing mH2A2 vs. H2B.

**Supplementary Table3**. GSEA analysis of Senescence and Autophagy in cancer WikiPathway in T-HEp3 cells overexpressing mH2A2 vs. H2B.

**Supplementary Table4**. Enrichr analysis of DEGs in T-HEp3 cells overexpressing mH2A2 vs. H2B.

**Supplementary Table5**. Lists of up- and down-regulated genes in D-HEp3 cells (vs. T-HEp3 cells) and in mH2A2-overexpressing cells (vs. H2B). The DEG data of D-HEp3 cells vs T-HEp3 cells is obtained in Gene Expression Omnibus (GSE181839).

**Supplementary Table6**. Lists of up- and down-regulated genes in OIS IMR90 cells (vs. growing IMR90 cells) and in mH2A2 overexpressing cells (vs. H2B). Senescent gene sets are obtained in Gene Expression Omnibus (GSE55949).

**Supplementary Table7**. Table of primer sequences.

**Supplementary Table8**. Table of antibodies and dilutions.

**Supplementary Table9**. Table of siRNAs, shRNAs information and guide RNA sequences for CRISPR-Cas9 knockout of human *MACROH2A1*.

## METHODS

### Cell lines & tumor growth studies

Tumorigenic (T-HEp3) HEp3 cells were derived from a lymph node metastasis from a HNSCC patient as described previously^42^. D-HEp3 cells were obtained by passing T-HEp3 cells for more than 40 generations *in vitro*^42^. When cultured *in vitro* all these cells were passaged in DMEM cell growth medium (Dulbeco’s modified medium with 10% of fetal bovine serum (FBS) and 100U penicillin/0.1 mg ml^-1^ streptomycin). MacroH2A overexpressing cells were generated by plKO-GFP-macroH2A encoding lentivirus infection of T-HEp3 cells then selected with puromycin (2.0 mg ml^-1^). Cells were routinely tested for mycoplasma (PCR Mycoplasma test kit PK-CA91-1096, PromoCell).

Tumour growth on chicken embryo chorioallantoic membranes (CAMs)^3^ or Balb/c nude mice (Charles River, #490) was performed as described previously^29^. All animal studies were approved by Institutional Animal Care and Use Committees (IACUC) at Mount Sinai School of Medicine Protocol ID: 08-0366. Briefly, 150×10^3^ ∼ 200 ×10^3^ T-HEp3 cells were inoculated on the CAM and allowed them to grow for the indicated times in each experiment (from 4 to 7 days). Dormant D-HEp3 cells were inoculated onto CAMs at 500×10^3^ cells per egg. For TGF-β2, (R&D systems, 302-B2) experiments on CAM, HEp3 cells were pre-treated *in vitro* for 24 h and then inoculated on CAMs for 6 days. CAM tumors were treated every day with TGF-β2 (10 ng ml^−1^). For siRNA experiments on CAM, HEp3 cells were pre-treated *in vitro* with scramble RNA or siRNA for 48hrs and then inoculated on CAMs for 5 days.

For HEp3 experimental metastasis assay, 3×10^5^ cells were tail-vein injected into BALB/c nu/nu female mice (4– 6 weeks). Mice were inspected every 48 hrs and were kept for 1 week or 3 weeks. At the end of week 1 or week 3, mice were sacrificed by euthanization, and lungs were collected.

For HEp3 spontaneous metastasis studies, 5×10^5^ cells were injected subcutaneously into BALB/c nu/nu female mice (4–6 weeks) in the interscapular region. Mice were inspected every 48 hrs and were kept (∼ 14 days) until primary tumours developed and grew up to >500 mm^3^. After surgical removal of the primary tumors, animals were randomly assigned to two groups. One as a negative control group was provided with regular drinking water, while the other group was provided with drinking water containing 2mg/ml doxycycline. Drinking water was replenished every 48hrs. Animals were monitored for local tumor recurrence every 48hrs for 6 weeks. At the end, mice were sacrificed by euthanization, and lungs were collected.

For NR2F1 agonist—C26 compound treatment, 3D organoid Assay, CAM tumor assay and spontaneous metastasis assay with C26 compound treatment were described previously^32^.

### Human specimens

Paraffin embedded sections from HNSCC primary tumors and matched lymph node metastasis lesions were obtained from the Cancer Biorepository at Icahn School of Medicine at Mount Sinai, New York, NY. Paraffin embedded tissue sections from lymph nodes biopsied from clinically diagnosed metastasis-free HNSCC patients were obtained from Dr. Christoph Sproll at the Heinrich-Heine-University Düsseldorf, Düsseldorf, Germany^40^. These patients presented small primary tumors, had no previously diagnosed malignancy in the head and neck region, and had not been subject to previous treatment. Samples were deidentified and obtained with institutional review board approval.

### Plasmids

Constitutively expressed human H2B-GFP and macroH2A2-GFP fusions were previously described^22^. Human macroH2A1.1 and macroH2A1.2-GFP fusions (transcript IDs XM_005272132.2 and NM_004893.3 respectively) were cloned into the pLKO.1 vector. Inducible macroH2A2-GFP constructs were generated as described^64^.

### Quantitative reverse transcriptome PCR (qPCR)

Total RNA was isolated from HEp3 cells using Trizol reagent (Invitrogen), and 2ug RNA was reverse-transcribed using M-MuLV reverse transcriptase enzyme (New England Biolabs #M0253S) following the manufacturer’s instructions. PCR was performed using PowerUP SYBR Green Master Mix (Applied Biosystems #A25741). Primers were purchased from IDT. Human tubulin was used as housekeeping gene control for all experiments. See Supplementary Table 6 for primer sequences.

### RNA interference

Gene-specific siRNAs (listed in Supplementary Table 9) were transfected to HEp3 cells at a final concentration of 40 nM using siPORT™ NeoFX™ Transfection Agent (Thermo Fisher #AM4510).

Lentiviral vectors of shRNAs were produced in HEK293T cells and viral supernatants were used to infect D-HEp3 cells. Detail information of shRNA generation was described previously^28^.

### Generation of macroH2A1 knockout cell lines

macroH2A1-knockout cell lines were generated using CRISPR/Cas9-targeted genome editing. Briefly, two separate macroH2A1 guide RNAs were designed and cloned into pLentiCRISPRv2^65^ (Addgene #52961). A non-targeting (NT) guide RNA was used as a control. Lentiviral vectors were produced in HEK293T cells and viral supernatants were used to infect D-HEp3 cells. Cells were then selected using puromycin (Sigma-Aldrich #P8833). macroH2A1 knockout was confirmed by western blot. See Supplementary Table 9 for guide RNA sequences.

### Immunofluorescence

Paraffin-embedded tissue sections were incubated in xylene (5 minutes twice) followed by graded ethanol rehydration (3 minutes each). Antigen retrieval for human primary tumours, lymph node metastasis tissues, mouse lung tissues and T-HEp3, D-HEp3 tumours was performed in 10 mM citrate buffer (pH 6) for 40 minutes using a steamer. Tissues were permeabilized in 0.1% TritonX-100 + PBS for 10 minutes. Tissue sections were blocked with 3% bovine serum albumin (BSA, Fisher Bioreagents) and 5% normal goat serum (NGS, Gibco #PCN5000) in PBS for 1 hour at room temperature. Primary antibodies (listed in Supplementary Table 8) in blocking buffer were incubated overnight at 4°C followed by washing and incubation with Alexa Fluor conjugated secondary antibodies (Invitrogen, 1:1000 dilution) at room temperature for 1-2 hour in the dark. Hoechst 33432 was used to stain nucleus (10 minutes) at room temperature. Slides were mounted with ProLong Gold Antifade reagent (Invitrogen #P36930). Images were obtained using Leica Software on a Leica SPE confocal microscope and analyzed using ImageJ software.

### RNA sequencing

Total RNA samples were prepared, poly A library construction and sequencing on Illumina NextSeq 500 at the ISMMS Sinai Genomics Core Facility, using the standard manufacture procedures. A detailed method of RNA sequencing and analysis was described previously^32^.

### RNA-Seq alignment and quality control

A total of 6 RNA-seq libraries (3 replicates each for overexpression and control) systems were processed using the same pipeline for compatibility. Quality control was performed using FastQC (v0.11.8)^66^. Trim Galore! (version 0.6.5 was used to trim the adapter sequences with a quality threshold of 20^67^. The human genome reference used was GRCh38.p13 and GENCODE release 36 was used as the transcriptome reference. The alignment is performed by using STAR aligner (v2.7.5b)^68^. Gene level read counts were obtained by using Salmon (v1.2.1) for all libraries^69^. All samples passed the quality control requirements with > 85% of reads uniquely mapping (>14 M uniquely mapped reads for each library) using STAR aligner.

### Differential expression and functional analysis

Differential expression analysis was performed using the gene level read counts and the DESeq2 (v1.28.1) R package^70^. Genes with less than 5 reads in total across all samples are filtered as inactive genes. A gene is considered differentially expressed if the adjusted p-value is less than 0.05 and the absolute fold change is greater than 0.5.

GSEA analysis with RNA sequencing data of T-HEp3 cells either overexpressing macroH2A2 or H2B tested the enrichment of ranked genes against molecular Signature Database Hallmark gene sets or Senescence and Autophagy in cancer in Wikipathway (See Supplementary Tables 2 and 6). Enrichr analysis of up- or down-regulated genes by macroH2A2 overexpression, respectively. The enrichment analysis is performed with Molecular Signature Database Hallmark gene sets, Kyoto Encyclopedia of Genes and Genomes (KEGG) Pathway Dataset and NCATS BioPlanet resource^71^ separately (See Supplementary Table 3).

### Principal component analysis and heatmaps

We performed the between sample normalization using the variance stabilizing transformation of the DESeq2 package. The 500 most variable genes are used to perform principal component analysis as well as calculating the Euclidean distances between each sample. Gene expression heatmaps show the z-scores of DESeq2 VST normalized gene-level read counts. The heatmaps were generated using all the DEGs identified to give an overview of the transcriptomic changes. All visualizations are generated using plotly R package (v4.9.2.1) except for heatmaps and volcano plots^72^. The heatmaps and volcano plots are generated using heatmaply (v1.1.0) and Glimma (v1.16.0) R packages respectively^73, 74^. R (v.4.0.3) is used to perform all bioinformatics analysis.

### Statistical analysis

All *in vitro* experiments were repeated at least three times unless indicated otherwise. For CAM tumor growth analysis, a minimum of 4 tumors were analyzed per group/experiment. For mouse experiments, a minimum of 5 mice per group were used for tumor growth, DCC and metastatic burden measurement and immunostaining analysis. Statistical analysis was performed on Prism software using unpaired t-test, Mann-Whitney test, and 2-way ANOVA with multiple comparison test. A p-value *≤* 0.05 was considered significant.

### Data availability

All datasets from this study have been uploaded to the Gene Expression Omnibus (GEO). RNA-seq dataset of T-HEp3 cells with macroH2A2 overexpression is accessible with the code of GSE182459. Previously published RNA-seq dataset of T-HEp3 vs. D-HEp3 cells and H3K27ac, H3K27me3 ChIP-seq in T-HEp3 and D-HEp3 cell lines are available under the accession codes GSE172115 and GSE181838. Previously published RNA-seq dataset of human fibroblasts with Oncogene-induced Senescence is available under the accession code GSE55949.

## REFERENCES

1. Risson, E., Nobre, A.R., Maguer-Satta, V. & Aguirre-Ghiso, J.A. The current paradigm and challenges ahead for the dormancy of disseminated tumor cells. Nature Cancer 1, 672–680 (2020).

2. Phan, T.G. & Croucher, P.I. The dormant cancer cell life cycle. Nat Rev Cancer 20, 398–411 (2020).

3. Aguirre Ghiso, J.A., Kovalski, K. & Ossowski, L. Tumor dormancy induced by downregulation of urokinase receptor in human carcinoma involves integrin and MAPK signaling. J Cell Biol 147, 89–104 (1999).

4. Crist, S.B. & Ghajar, C.M. When a House Is Not a Home: A Survey of Antimetastatic Niches and Potential Mechanisms of Disseminated Tumor Cell Suppression. Annu Rev Pathol 16, 409–432 (2021).

5. Sosa, M.S., Bernstein, E. & Aguirre-Ghiso, J.A. Epigenetic Regulation of Cancer Dormancy as a Plasticity Mechanism for Metastasis Initiation, in Tumor Dormancy and Recurrence. (eds. Y. Wang & F. Crea) 1–16 (Springer International Publishing, Cham; 2017).

6. Bragado, P. et al. TGF-beta2 dictates disseminated tumour cell fate in target organs through TGF-beta-RIII and p38alpha/beta signalling. Nat Cell Biol 15, 1351–1361 (2013).

7. Sosa, M.S. et al. NR2F1 controls tumour cell dormancy via SOX9-and RARbeta-driven quiescence programmes. Nat Commun 6, 6170 (2015).

8. Kobayashi, A. et al. Bone morphogenetic protein 7 in dormancy and metastasis of prostate cancer stem-like cells in bone. The Journal of experimental medicine 208, 2641–2655 (2011).

9. Ghajar, C.M. et al. The perivascular niche regulates breast tumour dormancy. Nature cell biology 15, 807–817 (2013).

10. Johnson, R.W. et al. Induction of LIFR confers a dormancy phenotype in breast cancer cells disseminated to the bone marrow.

11. Yumoto, K. et al. Axl is required for TGF-β2-induced dormancy of prostate cancer cells in the bone marrow. Scientific Reports 6, 36520 (2016).

12. Aguirre-Ghiso, J.A., Estrada, Y., Liu, D. & Ossowski, L. ERK(MAPK) activity as a determinant of tumor growth and dormancy; regulation by p38(SAPK). Cancer research 63, 1684–1695 (2003).

13. Aguirre-Ghiso, J.A., Liu, D., Mignatti, A., Kovalski, K. & Ossowski, L. Urokinase receptor and fibronectin regulate the ERK(MAPK) to p38(MAPK) activity ratios that determine carcinoma cell proliferation or dormancy in vivo. Molecular biology of the cell 12, 863–879 (2001).

14. Adam, A.P. et al. Computational identification of a p38SAPK-regulated transcription factor network required for tumor cell quiescence. Cancer research 69, 5664–5672 (2009).

15. Borgen, E. et al. NR2F1 stratifies dormant disseminated tumor cells in breast cancer patients. Breast Cancer Res 20, 120 (2018).

16. Zhang, R. et al. Formation of MacroH2A-containing senescence-associated heterochromatin foci and senescence driven by ASF1a and HIRA. Dev Cell 8, 19–30 (2005).

17. Chicas, A. et al. H3K4 demethylation by Jarid1a and Jarid1b contributes to retinoblastoma-mediated gene silencing during cellular senescence. Proceedings of the National Academy of Sciences 109, 8971–8976 (2012).

18. Salama, R., Sadaie M Fau - Hoare, M., Hoare M Fau - Narita, M. & Narita, M. Cellular senescence and its effector programs.

19. Ghiraldini, F.G., Filipescu, D. & Bernstein, E. Solid tumours hijack the histone variant network. Nat Rev Cancer 21, 257–275 (2021).

20. Sogabe, Y., Seno, H., Yamamoto, T. & Yamada, Y. Unveiling epigenetic regulation in cancer, aging, and rejuvenation with in vivo reprogramming technology. Cancer Sci 109, 2641–2650 (2018).

21. Costanzi, C. & Pehrson, J.R. Histone macroH2A1 is concentrated in the inactive X chromosome of female mammals. Nature 393, 599–601 (1998).

22. Gaspar-Maia, A. et al. MacroH2A histone variants act as a barrier upon reprogramming towards pluripotency. Nature Communications 4, 1565 (2013).

23. Barrero María J. et al. Macrohistone Variants Preserve Cell Identity by Preventing the Gain of H3K4me2 during Reprogramming to Pluripotency. Cell Reports 3, 1005–1011 (2013).

24. Pasque, V., Gillich, A., Garrett, N. & Gurdon, J.B. Histone variant macroH2A confers resistance to nuclear reprogramming. The EMBO Journal 30, 2373–2387 (2011).

25. Hu, W.H. et al. Loss of histone variant macroH2A2 expression associates with progression of anal neoplasm.

26. Sporn, J.C. & Jung, B. Differential regulation and predictive potential of MacroH2A1 isoforms in colon cancer.

27. Sporn, J.C. et al. Histone macroH2A isoforms predict the risk of lung cancer recurrence. Oncogene 28, 3423–3428 (2009).

28. Kapoor, A. et al. The histone variant macroH2A suppresses melanoma progression through regulation of CDK8. Nature 468, 1105–1109 (2010).

29. Aguirre-Ghiso, J.A., Ossowski, L. & Rosenbaum, S.K. Green fluorescent protein tagging of extracellular signal-regulated kinase and p38 pathways reveals novel dynamics of pathway activation during primary and metastatic growth. Cancer research 64, 7336–7345 (2004).

30. Ossowski, L. & Reich, E. Loss of malignancy during serial passage of human carcinoma in culture and discordance between malignancy and transformation parameters. Cancer research 40, 2310–2315 (1980).

31. Nobre, A.R. et al. Bone marrow NG2+/Nestin+ mesenchymal stem cells drive DTC dormancy via TGF-β2. Nature Cancer 2, 327–339 (2021).

32. Khalil, B.D. et al. An NR2F1-specific agonist suppresses metastasis by inducing cancer cell dormancy. Journal of Experimental Medicine 219 (2021).

33. Goddard, E.T., Bozic, I., Riddell, S.R. & Ghajar, C.M. Dormant tumour cells, their niches and the influence of immunity. Nat Cell Biol 20, 1240–1249 (2018).

34. Dasgupta, A., Lim, A.R. & Ghajar, C.M. Circulating and disseminated tumor cells: harbingers or initiators of metastasisã Mol Oncol 11, 40–61 (2017).

35. Naume, B. et al. Clinical outcome with correlation to disseminated tumor cell (DTC) status after DTC-guided secondary adjuvant treatment with docetaxel in early breast cancer. J Clin Oncol 32, 3848–3857 (2014).

36. Fluegen, G. et al. Phenotypic heterogeneity of disseminated tumour cells is preset by primary tumour hypoxic microenvironments. Nat Cell Biol 19, 120–132 (2017).

37. Ossowski, L., Russo-Payne, H. & Wilson, E.L. Inhibition of urokinase-type plasminogen activator by antibodies: the effect on dissemination of a human tumor in the nude mouse. Cancer research 51, 274–281 (1991).

38. Ossowski, L., Russo, H., Gartner, M. & Wilson, E.L. Growth of a human carcinoma (HEp3) in nude mice: rapid and efficient metastasis. J Cell Physiol 133, 288–296 (1987).

39. Sproll, C., Fluegen, G. & Stoecklein, N.H. Minimal Residual Disease in Head and Neck Cancer and Esophageal Cancer. Adv Exp Med Biol 1100, 55–82 (2018).

40. Sproll, C. et al. Immunohistochemical detection of lymph node-DTCs in patients with node-negative HNSCC. Int J Cancer 140, 2112–2124 (2017).

41. Chery, L. et al. Characterization of single disseminated prostate cancer cells reveals tumor cell heterogeneity and identifies dormancy associated pathways. Oncotarget 5, 9939–9951 (2014).

42. Ossowski, L. & Reich, E. Changes in malignant phenotype of a human carcinoma conditioned by growth environment. Cell 33, 323–333 (1983).

43. Kuleshov, M.V. et al. Enrichr: a comprehensive gene set enrichment analysis web server 2016 update.

44. Lee, H.J. et al. Proteomic and Metabolomic Characterization of a Mammalian Cellular Transition from Quiescence to Proliferation. Cell Rep 20, 721–736 (2017).

45. Honma, S. et al. Dec1 and Dec2 are regulators of the mammalian molecular clock. Nature 419, 841–844 (2002).

46. Sancar, A. & Van Gelder, R.N. Clocks, cancer, and chronochemotherapy. Science 371 (2021).

47. Acosta, J.C. et al. A complex secretory program orchestrated by the inflammasome controls paracrine senescence. Nat Cell Biol 15, 978–990 (2013).

48. Duarte, L.F. et al. Histone H3.3 and its proteolytically processed form drive a cellular senescence programme. Nat Commun 5, 5210 (2014).

49. Nagy, A., Munkacsy, G. & Gyorffy, B. Pancancer survival analysis of cancer hallmark genes. Sci Rep 11, 6047 (2021).

50. Ruth, J.R. et al. Cellular dormancy in minimal residual disease following targeted therapy. Breast Cancer Res 23, 63 (2021).

51. Shachaf, C.M. et al. MYC inactivation uncovers pluripotent differentiation and tumour dormancy in hepatocellular cancer. Nature 431, 1112–1117 (2004).

52. Redelsperger, I.M. et al. Stability of Doxycycline in Feed and Water and Minimal Effective Doses in Tetracycline-Inducible Systems. J Am Assoc Lab Anim Sci 55, 467–474 (2016).

53. Wilson, A. et al. Hematopoietic stem cells reversibly switch from dormancy to self-renewal during homeostasis and repair. Cell 135, 1118–1129 (2008).

54. Chung, C.Y. et al. Cbx8 Acts Non-canonically with Wdr5 to Promote Mammary Tumorigenesis. Cell Rep 16, 472–486 (2016).

55. Bragado, P. et al. Analysis of marker-defined HNSCC subpopulations reveals a dynamic regulation of tumor initiating properties. PloS one 7, e29974 (2012).

56. Borgen, E. et al. NR2F1 stratifies dormant disseminated tumor cells in breast cancer patients. Breast Cancer Research 20, 1–13 (2018).

57. Singh, D.K. et al. Epigenetic reprogramming of DCCs into dormancy suppresses metastasis <em>via</em> restored TGFβ–SMAD4 signaling. bioRxiv, 2021.2008.2001.454684 (2021).

58. Timinszky, G. et al. A macrodomain-containing histone rearranges chromatin upon sensing PARP1 activation.

59. Kustatscher, G., Hothorn M Fau -Pugieux, C., Pugieux C Fau - Scheffzek, K., Scheffzek K Fau - Ladurner, A.G. & Ladurner, A.G. Splicing regulates NAD metabolite binding to histone macroH2A.

60. Posavec Marjanović, M. et al. MacroH2A1.1 regulates mitochondrial respiration by limiting nuclear NAD+ consumption. Nature Structural & Molecular Biology 24, 902–910 (2017).

61. Chen, H. et al. MacroH2A1 and ATM Play Opposing Roles in Paracrine Senescence and the Senescence-Associated Secretory Phenotype.

62. Hsu, C.-J., Meers, O., Buschbeck, M. & Heidel, F.H. The Role of MacroH2A Histone Variants in Cancer. Cancers 13 (2021).

63. Lujambio, A. et al. Non-cell-autonomous tumor suppression by p53. Cell 153, 449–460 (2013).

64. Sun, Z. et al. Transcription-associated histone pruning demarcates macroH2A chromatin domains. Nature Structural & Molecular Biology 25, 958–970 (2018).

65. Sanjana, N.E., Shalem, O. & Zhang, F. Improved vectors and genome-wide libraries for CRISPR screening. Nature Methods 11, 783–784 (2014).

66. Andrews, S. (Babraham Bioinformatics; 2010).

67. Krueger, F. (Babraham Bioinformatics, Babraham Institute, Cambridge, United Kingdom. ; 2012).

68. Dobin, A. et al. STAR: ultrafast universal RNA-seq aligner.

69. Patro, R., Duggal, G., Love, M.I., Irizarry, R.A. & Kingsford, C. Salmon provides fast and bias-aware quantification of transcript expression.

70. Love Mi Fau - Huber, W., Huber W Fau - Anders, S. & Anders, S. Moderated estimation of fold change and dispersion for RNA-seq data with DESeq2.

71. Huang, R. et al. The NCATS BioPlanet – An Integrated Platform for Exploring the Universe of Cellular Signaling Pathways for Toxicology, Systems Biology, and Chemical Genomics. Frontiers in Pharmacology 10, 445 (2019).

72. C, S. (Chapman and Hall, 2020).

73. Galili, T., O’Callaghan, A., Sidi, J. & Sievert, C. heatmaply: an R package for creating interactive cluster heatmaps for online publishing.

74. Su, S. et al. Glimma: interactive graphics for gene expression analysis.

