## Supplementary figures for "MacroH2A impedes metastatic growth by enforcing a discrete dormancy program in disseminated cancer cells"

### Supplementary Figure 1.

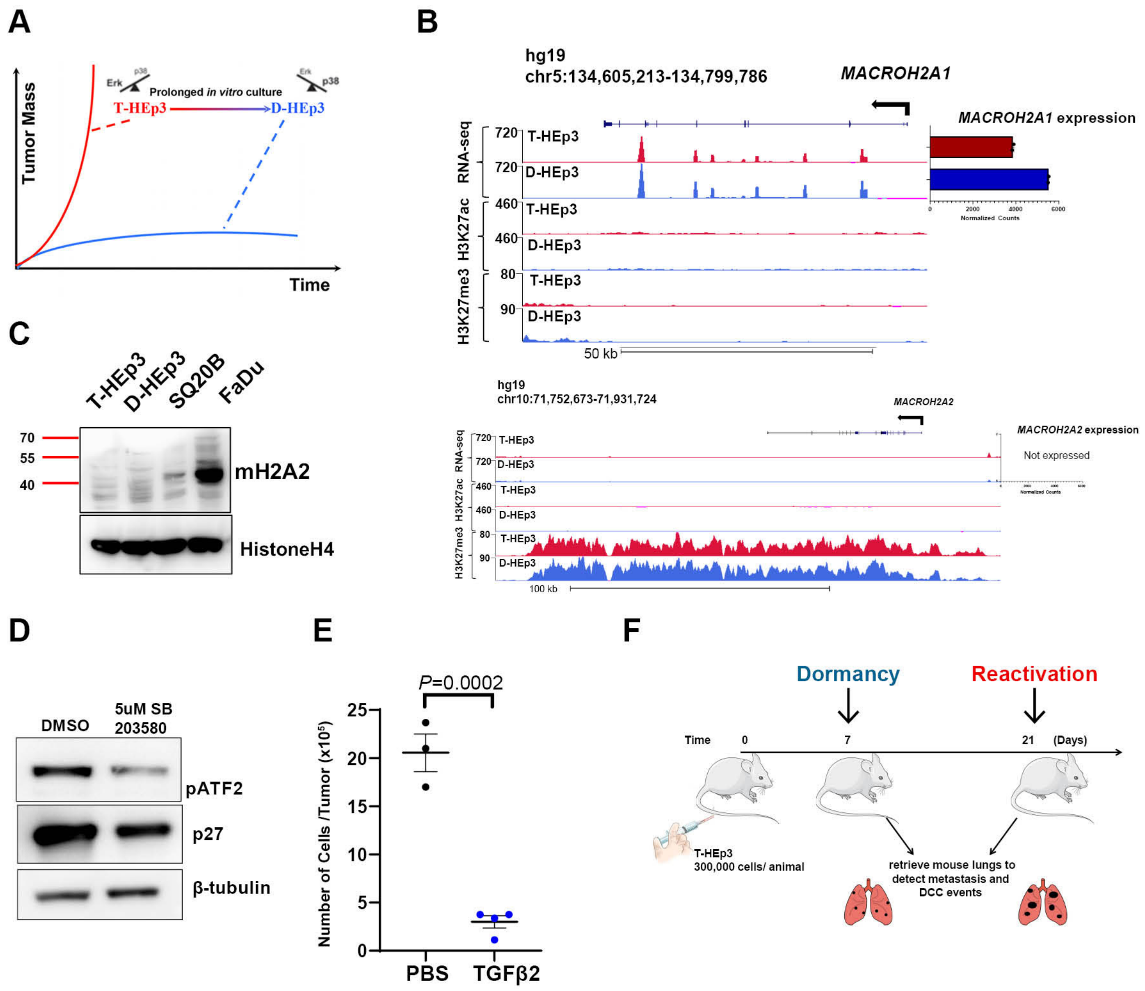

### Supplementary Fig.2

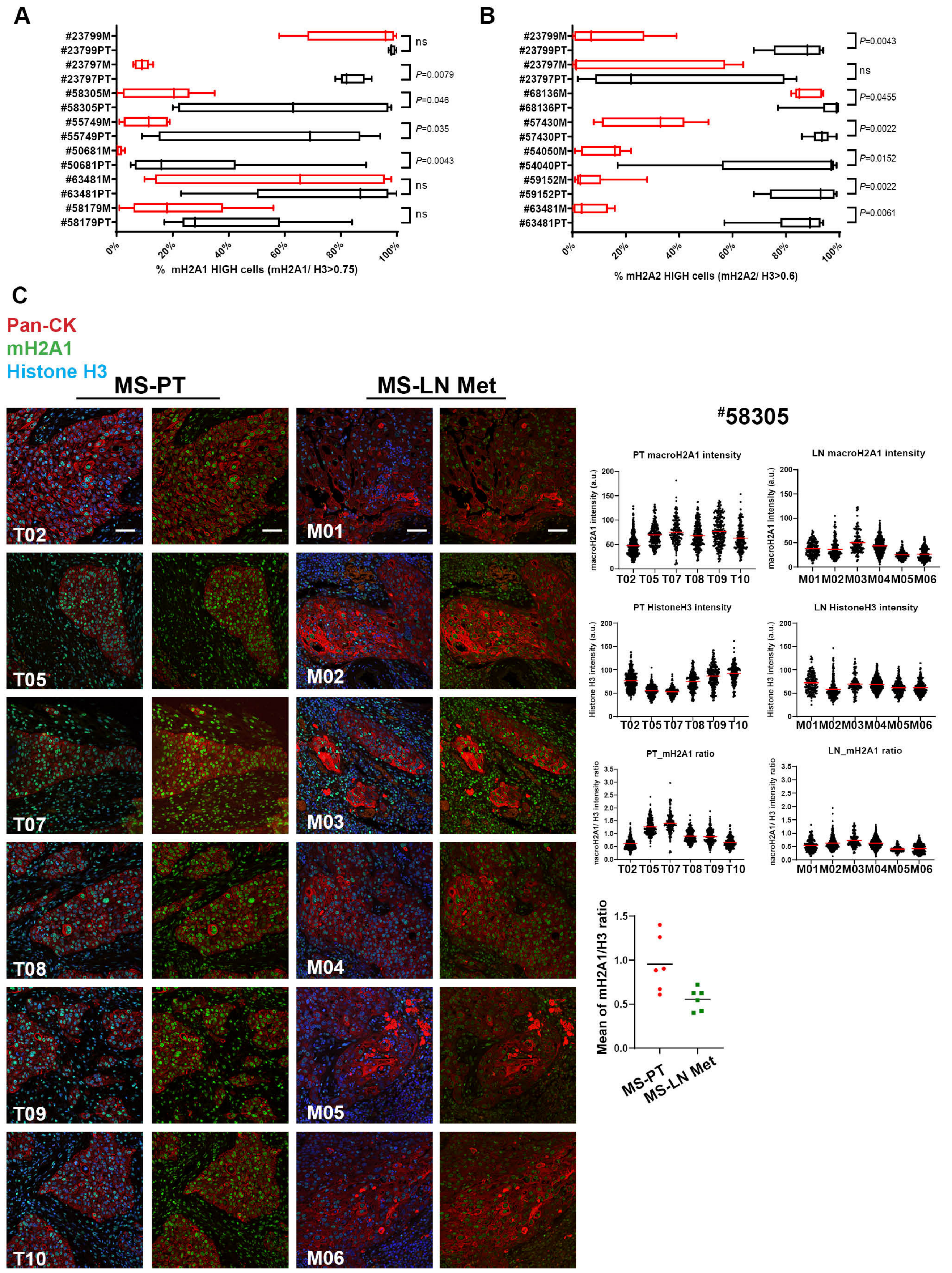

### Supplementary Fig.2

D

Pan-CK
mH2A2
Histone H3

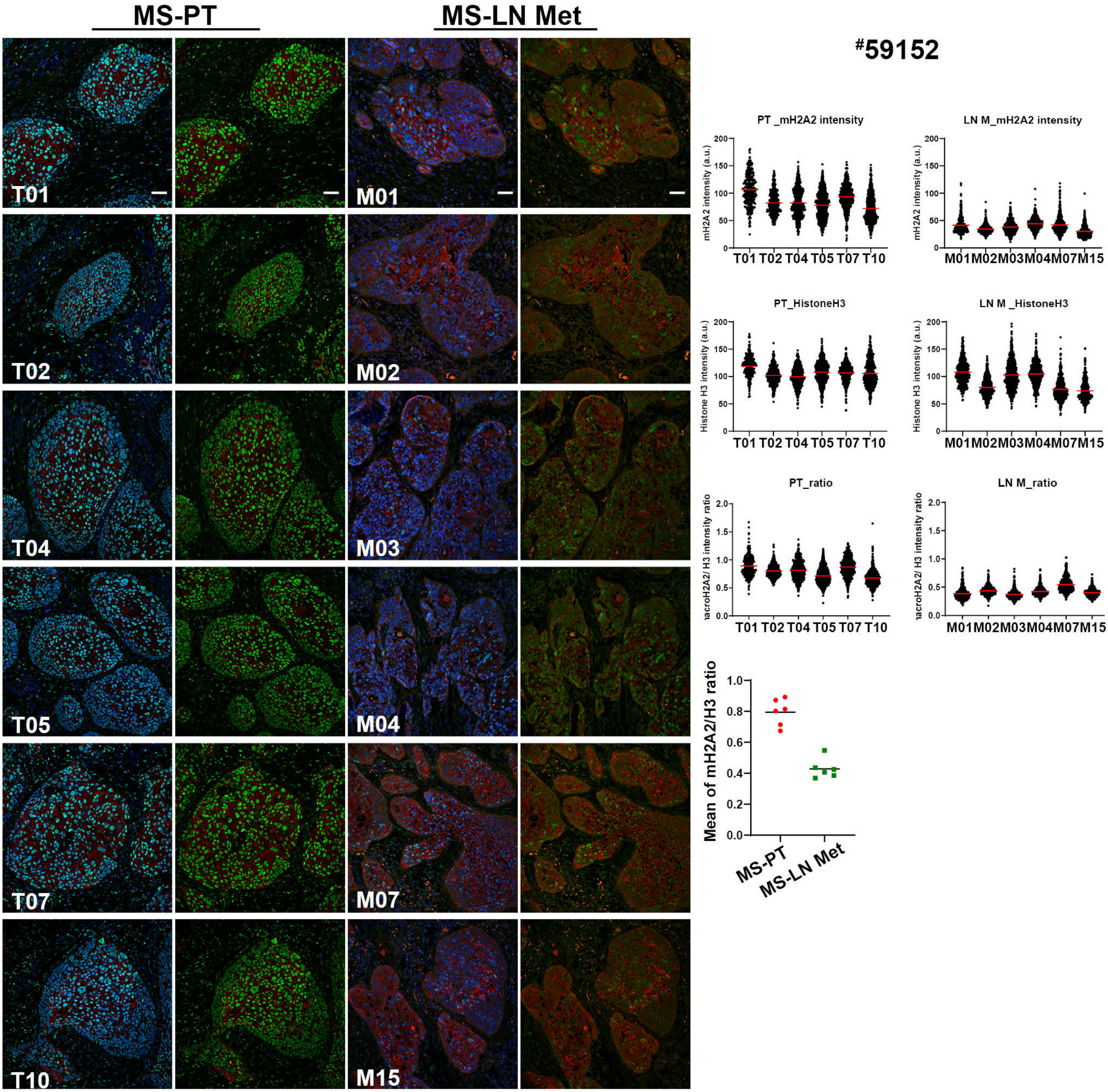


E

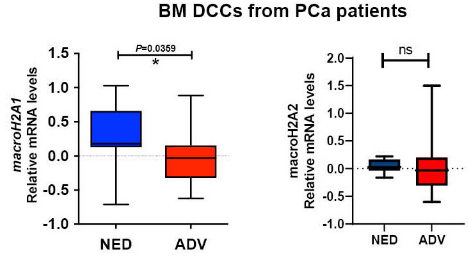

### Supplementary Fig. 3

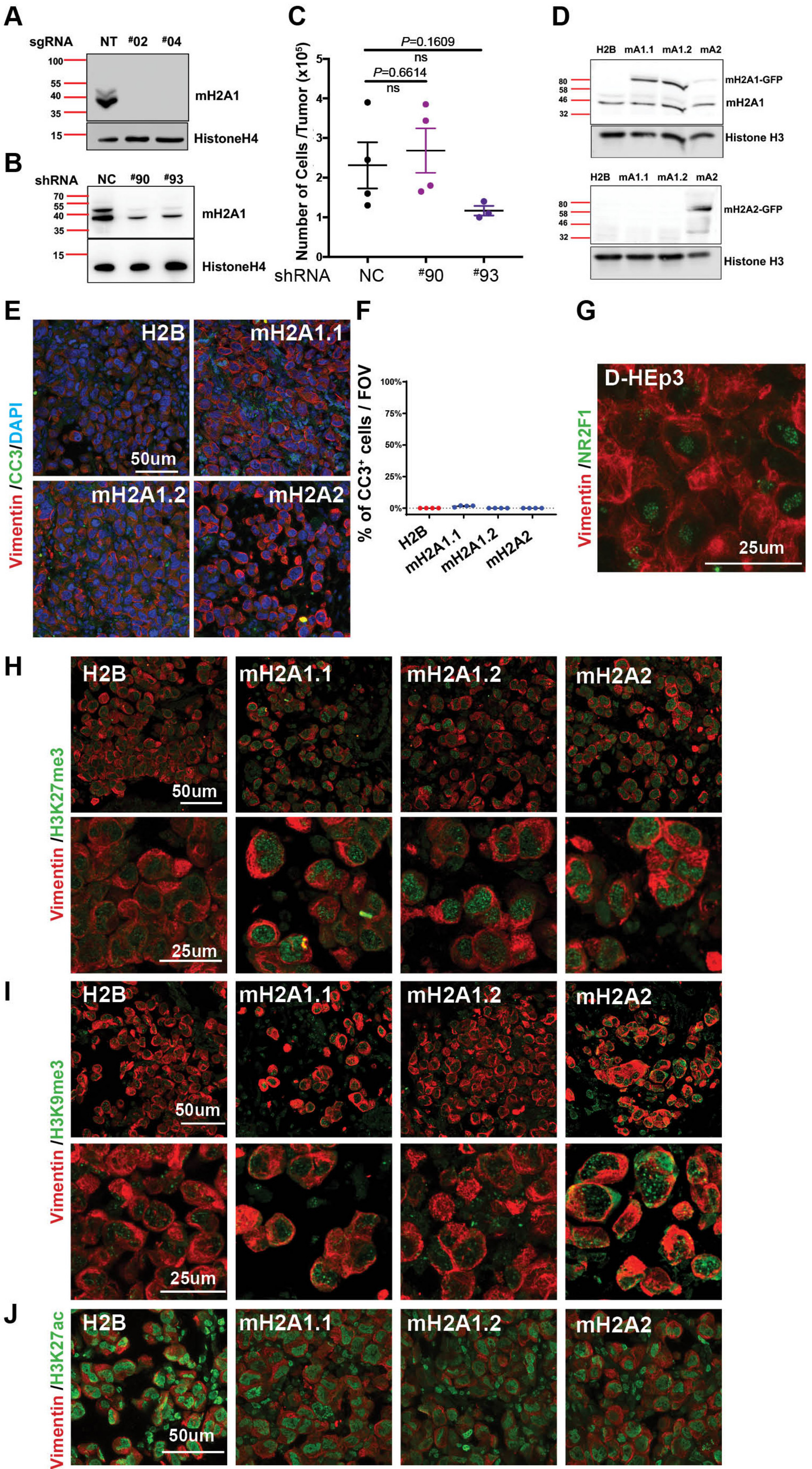

### Supplementary Fig.4

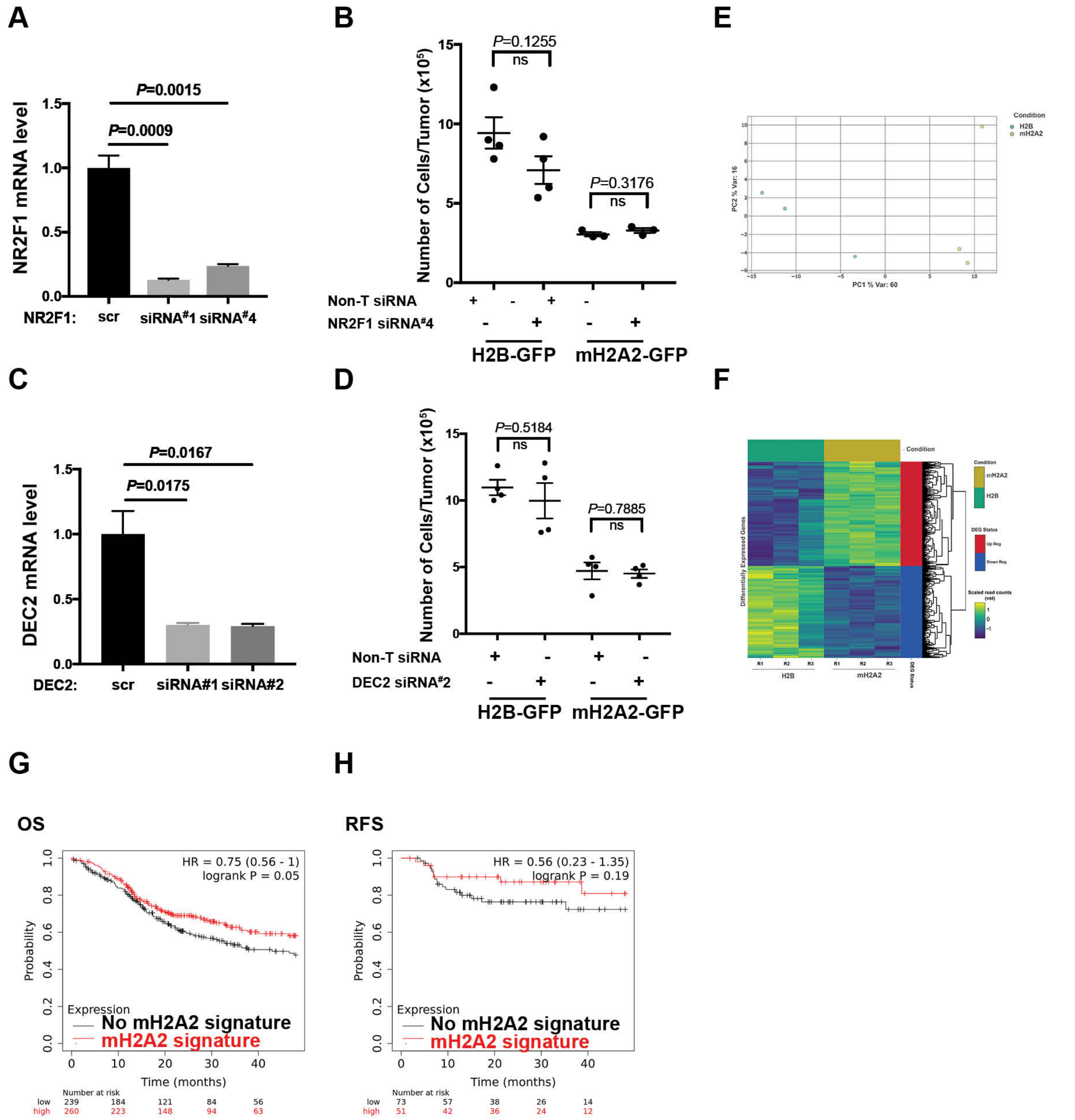

### Supplementary Fig.5

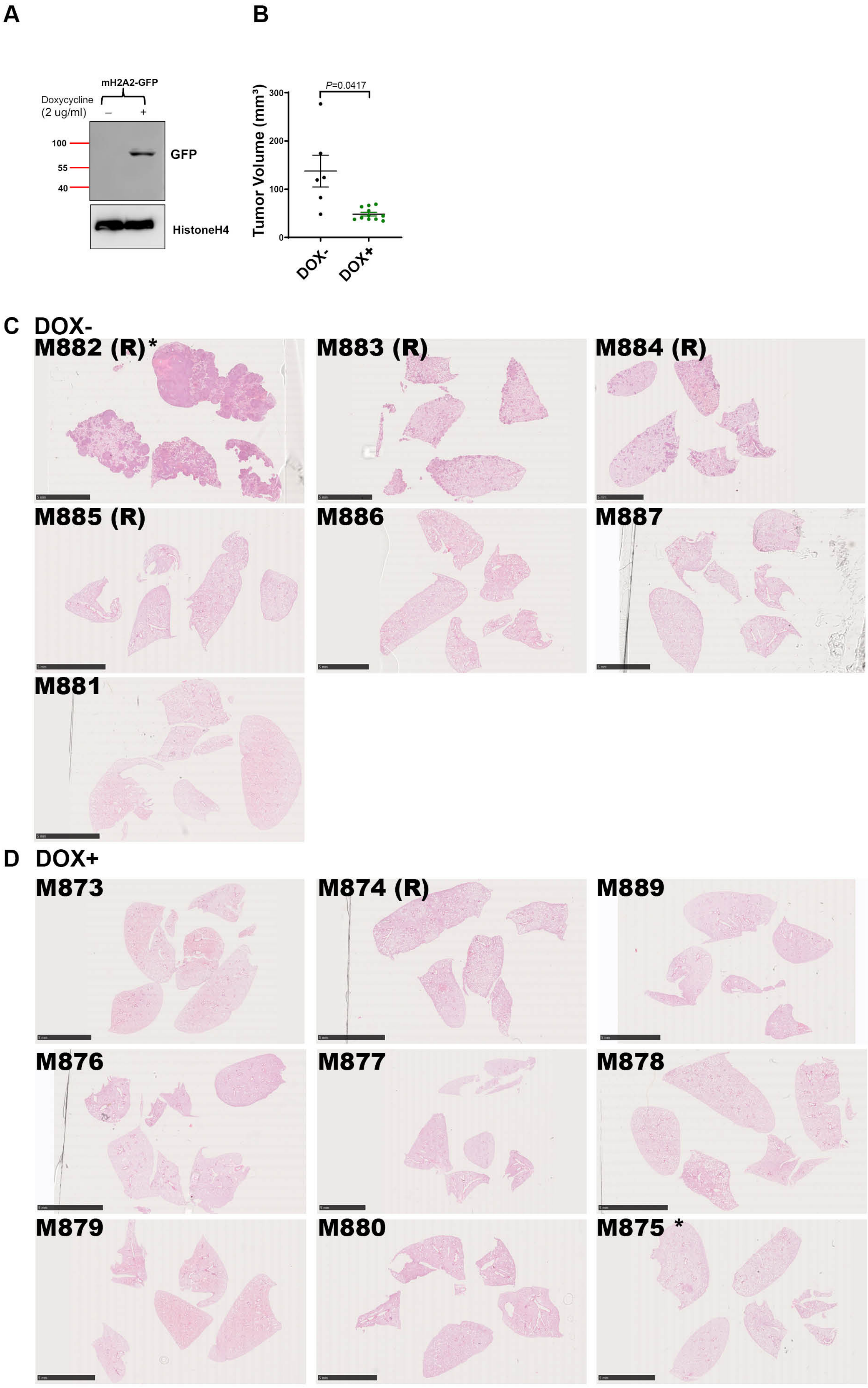
